# Interpersonal comparison of utility by measuring neural activity

**DOI:** 10.1101/2021.06.04.447048

**Authors:** Kaosu Matsumori, Kazuki Iijima, Yukihito Yomogida, Kenji Matsumoto

## Abstract

Aggregating well-being across individuals to reach collective decisions is one of the most fundamental problems in our society. Interpersonal comparisons of utility are pivotal and inevitable for well-being aggregation, because if utility is not interpersonally comparable, there is no rational aggregation procedure that simultaneously satisfies even mild conditions for validity (Arrow’s impossibility theorem); however, there are as yet no scientific methods for interpersonal comparison of utility. In this study, we developed a method based on brain signals. We found that the anterior cingulate and its adjacent ventromedial prefrontal activity is correlated with changes in expected utility. The ratio of lower- and higher-income participants’ neural signals coincided with estimates of their psychological pleasure by “impartial spectators.” We confirmed the validity of our interpersonal utility comparison method using an independent large-scale dataset and used the aggregated well-being from our experimental data to derive an optimal decision rule. These findings suggest that our interpersonal comparison method enables scientifically reasonable well-being aggregation by escaping Arrow’s impossibility and has implications for the fair distribution of economic goods. Our method can be used for evidence-based policy-making in nations that use cost-benefit analyses or optimal taxation theory for policy evaluation.

## Introduction

Consider a situation where you decide to give a piece of candy to one of two people. If one person wants it and the other does not, it is easy to make your decision. What if both people say, “I want it”? If you want to give the candy to the person who will derive more pleasure from receiving it, you need to be able to compare the potential pleasure the two people would derive from having the candy. You may be able to make a decision if you know the two people well; however, it is extremely difficult to make a decision with enough objectivity to convince a person with a different opinion about the two people’s potential enjoyment of the candy.

This situation illustrates a serious problem in modern societies that must consider taxation and wealth distribution policies, because the utilities of large numbers of unknown people, ranging from the very poor to the very rich, have to be compared. This “interpersonal comparison of utility” is pivotal and inevitable for policy evaluation, but it is almost completely ignored in current economics, because interpersonal comparisons of utility have been regarded as having no empirical meaning (Arrow 1963; Robbins 1932, 1938; Sen 2018). This stems from the history of utility concepts.

Bentham’s original sense of utility (psychological utility) was psychological pleasure (Bentham 1789). Since psychological utility cannot be directly measured from behavior, a new concept of utility (decision utility), which can be operationally defined by the agent’s choice behaviors, emerged as a surrogate for psychological utility at the beginning of the 20th century (Kreps 2012; Moscati 2018) (Supplementary Information Section 1).

Among current views of decision utility, “expected utility” is the most informative (Von Neumann and Morgenstern 2007) (Figure 1A–C). Expected utility can be expressed on a scale of 0 for the worst option to 1 for the best option (0–1 rescaling) without losing any information (Supplementary Information Section 1). This characteristic indicates that expected utility (and other decision utilities) is by definition only *intra*personally and not *inter*personally comparable (Figure 1E, F).

**Figure 1.**
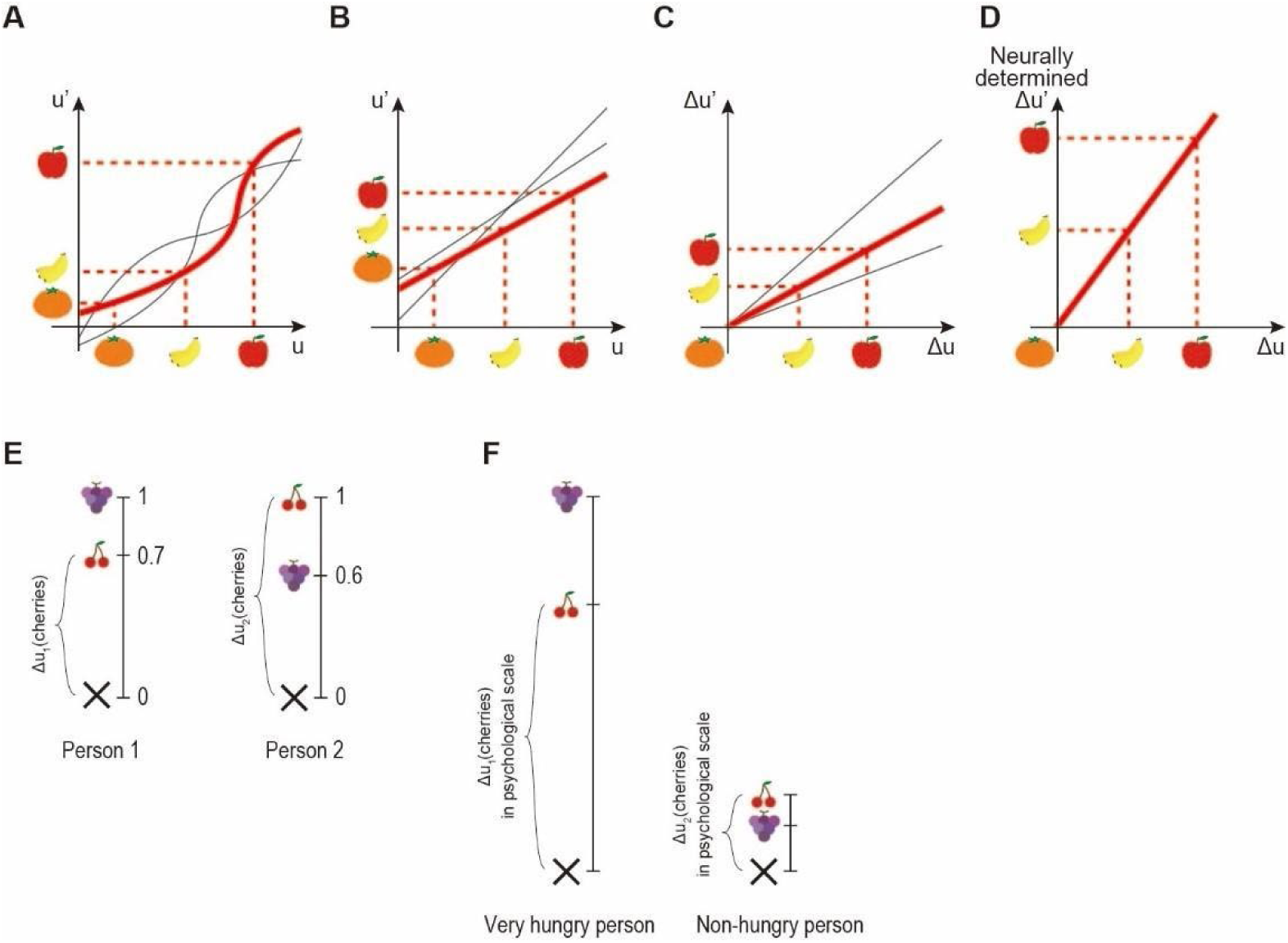
Concepts of utility. (**A**) Utility as an ordinal scale (ordinal utility). Each axis represents the preference for an apple over a banana and a banana over an orange. Only the order of the options is meaningful in this scale, so any monotonically increasing transformation is permitted without losing any information. A representative (red line) and two variants (thin gray lines) are shown. (**B**) Utility as an interval scale (cardinal utility). Each axis represents utilities in which the utility difference between an apple and a banana is the same as the utility difference between a banana and an orange. Only the distances between the options are meaningful in this scale, so any positive linear transformation (u’ = αu + β, α > 0) is permitted without losing any information. A representative (red line) and two variants (thin gray lines) are shown. (**C**) Utility difference as a ratio scale. If the cardinal utility is given, the utility difference (Δu) is a ratio scale by subtracting the reference option (an orange in the figure). Any positive scalar multiplication (Δu’ = αΔu, α > 0) is permitted without losing any information. (**D**) Given a neuroscientific measurement method, a single scale for Δu can be determined by using the axis of neural activity correlating with Δu in each individual. (**E**) As the lengths of the scales for cardinal utility have no meaning, a 0–1 rescaling is applied to express the utilities of two individuals. Δu_1_(cherries) = 0.7 and Δu_2_(cherries) = 1, where Δu(⋅) = u(⋅) - u(none). Thus, Δu_1_(cherries) < Δu_2_(cherries), although this comparison has no empirical meaning. (**F**) A modified case of **E**. Suppose Person 1 is very hungry and Person 2 is not hungry at all, we intuitively recognize that a hungry person derives more pleasure from receiving cherries than a person who is not hungry. Thus, we can give a specific meaning to the lengths of the scales to match our intuition by making the hungry person’s scale longer and the non-hungry person’s scale shorter, thus providing a larger psychological utility of cherries for the hungry person than for the non-hungry person in contrast to **E**.

Since the concept of decision utility was popularized, interpersonal comparisons of utility have been regarded as neither objective nor scientific (Robbins 1932, 1938) for nearly a century. This is a serious problem caused by replacing psychological utility with decision utility. Using decision utility alone, the utility for a slice of bread cannot be compared even between a starving person and a billionaire (Sen 2018).

Analyses based on decision utility show that a competitive equilibrium based on an ideal market can realize a Pareto efficient state; in this state, no one’s utility can increase any further without decreasing another’s utility (Supplementary Figure 1); however, Pareto efficiency is not considered a sufficient condition for proper resource allocation, because Pareto efficient resource allocation does not exclude highly unequal states. The next question is, “How can we reach an appropriate resource allocation method that satisfies conditions beyond Pareto efficiency?” Kenneth Arrow denied the existence of such a method by proving mathematically—Arrow’s impossibility theorem—that there is no rational procedure that simultaneously satisfies even very mild conditions for validity without interpersonally comparable utility (Arrow 1963; Sen 2009, 2018). Nevertheless, governments are often compelled to evaluate policies to make collective decisions in which interpersonal comparisons of utility are required. Because there is no appropriate method to compare utilities interpersonally, policy evaluations are often conducted by assuming that the utility of a penny is exactly the same across all income levels—social cost-benefit analysis—from the poor to the very rich (Boardman et al. 2018; HM Treasury 2018). This assumption is unrealistic and unintuitive (Boardman et al. 2018; Sen 2018). If we could find an appropriate method for interpersonal comparisons of utility, we could deal with the distribution problem properly.

To escape from Arrow’s impossibility, it is essential to extend the informational basis of policy evaluation by developing an appropriate interpersonal comparison method. An appropriate method for interpersonal comparison of utility has to satisfy the following two conditions. First, the interpersonal comparison method must rest on scientific demonstration rather than on a mere stipulation of ethical principles (Robbins 1932, 1938). This condition warrants the objectivity of the results of interpersonal comparisons of utility. Second, the interpersonal comparison method must effectively capture human intuitions about utility and be validated by their coherence, as was done when the temperature scale based on the fluid expansion was adopted in its initial stages (Chang 2004) (Supplementary Information Section 1). Satisfying this condition must serve as a starting point for subsequent scientific inquiry (Chang 2004). One popular idea among both economists and philosophers is that, to solve the problem of interpersonal comparisons of utility, we need to look at how ordinary people make such comparisons in everyday life (Harsanyi 1955; Rossi 2014), because ordinary people appear to make interpersonal comparisons of utility with relative ease and apparent success (Davidson 2004; Rossi 2014).

In this study, we developed a scientific method for interpersonal comparisons of utility based on brain-derived signals from functional magnetic resonance imaging (fMRI). In addition, we validated the interpersonal comparison method by comparing utility based on MRI signals with impartial spectators’ subjective estimations. To further establish the external validity of our findings, we analyzed data from the Adolescent Brain Cognitive Development (ABCD) study, involving over 5,000 children. Finally, we applied the interpersonal comparison method to an actual distribution problem.

## Results

### Neural representation of utility

We focused on a neural representation of utility to construct a method for interpersonal comparison of utility for the following reasons. First, any economic choice behavior that reveals decision utility consists of voluntary muscle movements generated by signals that originate in the brain. Second, it is natural to assume that economic choice behavior that reveals decision utility is determined by psychological utility. Furthermore, as mental states—including psychological utility—are realized by brain states, interpersonally comparable utility may be found in the activities of the responsible brain areas.

In the past quarter century, neuroscience studies have uncovered neural circuits that represent subjective values, known as the “value system” (Bartra, McGuire, and Kable 2013; Glimcher and Fehr 2013) (Supplementary Information Section 2). The value system consists of a set of brain regions, including the medial prefrontal cortices and anterior striatum, that learn the values of choice options by receiving inputs from the midbrain dopaminergic nuclei that encode prediction error of decision utility (Schultz et al. 1997; Stauffer et al. 2014; Sutton and Barto 2018) (i.e., the difference between the utility of an obtained outcome and the expected utility of a lottery; Δu).

To examine the neural representation of utility, we identified a representation that correlates with the prediction error of decision utility (Δu), which commonly works irrespective of reward type (i.e., food and money), rather than the prediction error of reward amount. In our gambling experiment, we engaged 63 participants in two types of gambling tasks: a food gambling task and a monetary gambling task. We used brain imaging data from 56 and 60 participants during the food gambling task and the monetary gambling task, respectively, after excluding data from 7 and 3 participants, respectively, due to head movements during the scan, insufficient motivation to obtain the food reward, and a technical problem in data storage (see *Methods*). Because intrapersonal utility function for moderate amounts of money is approximately linear (Abdellaoui et al. 2008; Booij and Kuilen 2009; Wakker and Deneffe 1996), it was expected that the prediction error of utility and the prediction error of reward amount will be strongly correlated in the monetary gambling task. Therefore, we used the food gambling task to identify brain regions that correlate with the prediction error of decision utility (Δu) rather than that of the reward amount.

In the food gambling task, participants chose between a sure payoff of food (snack) tickets and a lottery entailing a 50/50 chance of gaining one of two quantities of food tickets (Figure 2A). We determined each participant’s 0–1 scaled utility function for food tickets using the fractile method, with considering probability weighting function (Stauffer et al. 2014) (see *Methods*, Supplementary Figure 2). The slope of the utility function for most of the participants decreased as the amount of food increased (Figure 2B), indicating that their marginal utilities diminished in the range 1–300 food tickets.

**Figure 2.**
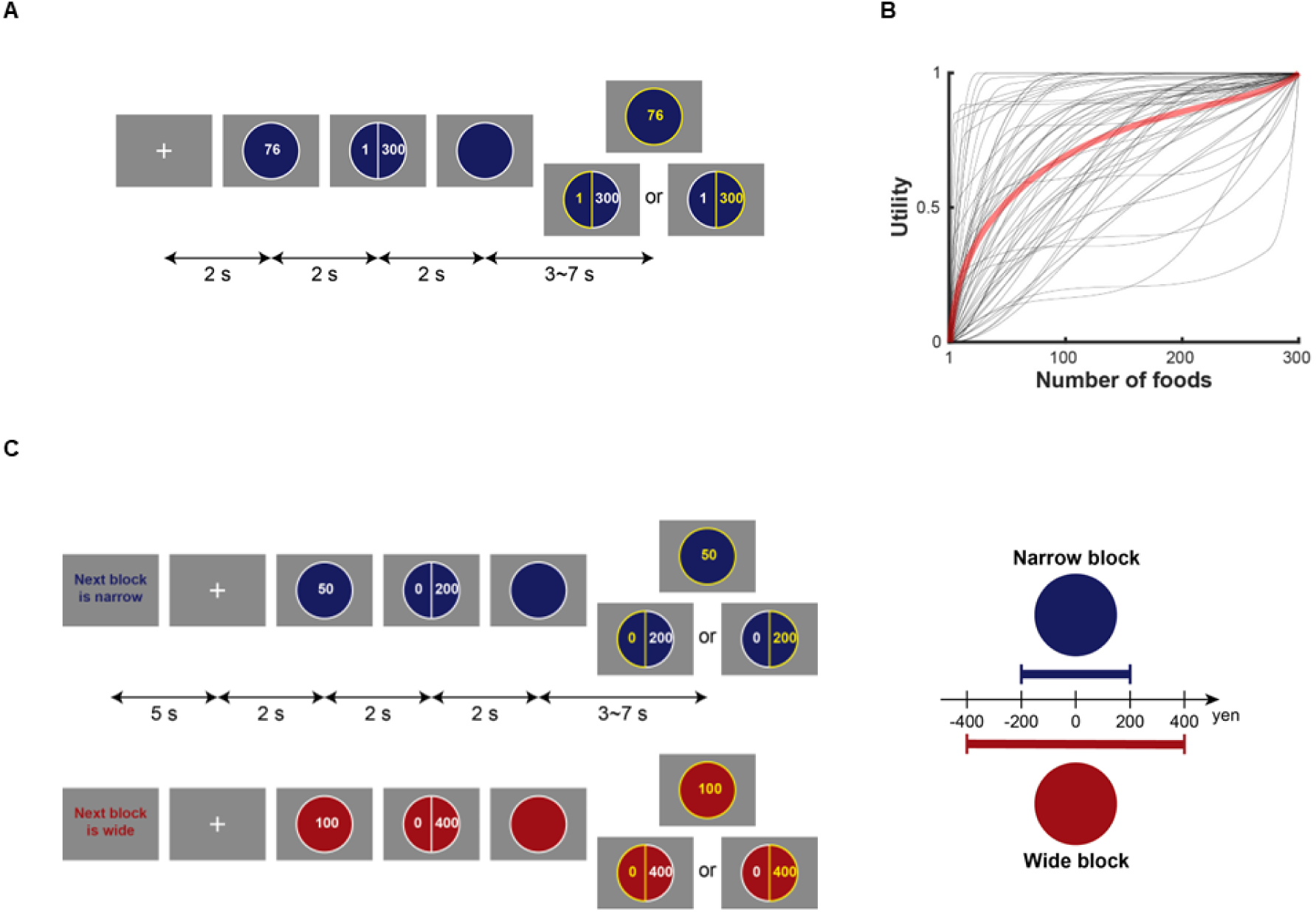
Gambling tasks. (**A**) Food gambling task. Participants chose between a sure (riskless) offer and a lottery to get indicated amounts of snacks as reward. (**B**) The experimentally obtained utility functions for food amounts. 0–1 rescaling was applied to the utility functions. The utility curves of the participants (thin black line) and their averaged curve (thick red line) are shown. (**C**) Monetary gambling task. Participants received money as a reward. This task consisted of alternating narrow and wide blocks with corresponding offer ranges (-¥200 - +¥200 and -¥400 - +¥400) as illustrated in the panel on the right.

Since we included trials that offered a random number of food tickets, although these trials were not used for the fractile method in the food gambling task (see *Methods*), we could use them to validate the shape of each participant’s utility function. The percentage of trials in which participants chose the offer with larger estimated utility was 77.13% on average (SD 10.28%) and the choice behaviors were, after consideration of the probability weighting, more accurately predicted by expected utility than expected reward amount (t(55) = 4.58, *P* < 0.001 two-sided), suggesting that the utility functions were effective in accounting for the participants’ behaviors.

Next, we analyzed the fMRI data to identify the brain regions whose activation correlated with utility. As we had an a priori hypothesis on the activation of the value system (Bartra et al. 2013; Glimcher and Fehr 2013), the analysis was performed for voxels within the explicit anatomical mask that covers the medial prefrontal cortices and anterior striatum. We found that the activation of a brain region consisting of the anterior cingulate cortex and its adjacent ventromedial prefrontal cortex (ACC/vmPFC) (peaked at [0, 38, -4] MNI coordination), correlated with the prediction error of utility (Δu) (GLM1) (red and yellow areas in Figure 3A (t(55) = 4.79, FWE corrected *P* = 0.008 one-sided). Importantly, the activity of this region, as well as others, did not correlate with the prediction error of the number of food tickets (GLM2). This indicates that activity in this region correlates with the prediction error of utility rather than that of the reward amount per se.

**Figure 3.**
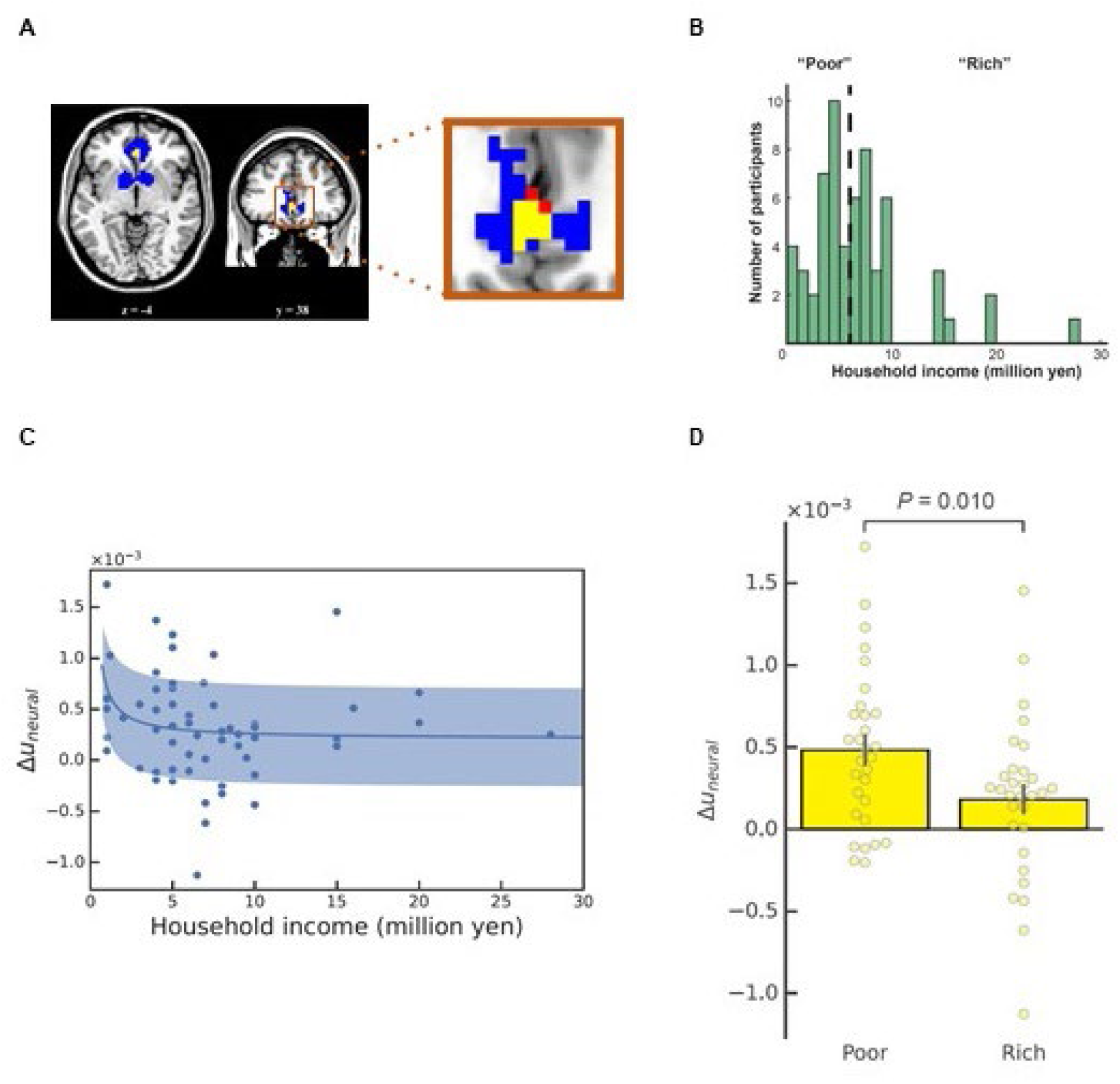
Interpersonal comparison of neural measures for utility. (**A**) Neural representations of utility prediction error. Activated regions in the brain are shown for the food gambling task (red), monetary gambling task (blue), and their overlapped region (yellow) (*P* < 0.05, FWE corrected). The area in the frame is magnified at right. The overlapped region in the anterior cingulate cortex and its adjacent ventromedial prefrontal cortex (ACC/vmPFC) is called the “utility region”. The PSC in the utility region per 1 yen is defined as Δu_neural_. (**B**) Distribution of annual household income for the participants in the gambling tasks. (**C**) Δu_neural_ of the participants are plotted against their household income. Δu_neural_ was inversely proportional to income (β_income-1_ = 0.247 ± 0.132 (mean ± SE), t(56) = 1.874, *P* = 0.033 one-sided) after regressing out age and sex. The shaded area indicates SD. (**D**) When the participants were split into a “poor” and a “rich” group (indicated by the vertical dashed line at 6 million yen in **B**), the poor group showed significantly greater Δu_neural_ than the rich group (t(58) = 2.39, *P* = 0.010 one-sided, BF_+0_ = 5.723). Error bars indicate SEM across group members.

Next, we examined whether the activity of the ACC/vmPFC region that correlated with utility prediction errors in the food gambling task also correlated with utility prediction errors in the monetary gambling task. In the monetary gambling task, participants chose between a guaranteed amount of money and a lottery for money with a 50% payoff probability in two different conditions: a narrow block that had a narrow range of reward amounts (¥-200∼200) and a wide block that had a wide range of reward amounts (¥- 400∼400) (1 USD ≈ 150 JPY (¥)) (Figure 2C). This allowed us to examine whether brain activity was influenced by different ranges of reward amounts. We compared the percent signal changes (PSC) of the ACC/vmPFC region evoked by the utility prediction errors per ¥1 between the narrow and wide block using both classical hypothesis testing and Bayesian hypothesis testing based on the Bayes factor (BF). We found moderate evidence indicating the absence of an influence across different ranges of reward amounts (t(59) = 0.36, *P* = 0.360 one-sided, BF_+0_ = 0.193, median posterior δ = 0.044 with 95% CI [-0.201, 0.290]) (GLM3) (Supplementary Figure 3). We ran another GLM (GLM4) and identified the voxels representing utility prediction errors in our anatomical mask without discriminating between the narrow and the wide block (peaked at [-9, 5, -10] and [9, 8, -7] (t(59) = 9.05, t(59) = 7.68) in the striatum and at [-6, 38, -4] (t(59) = 6.57) in the vmPFC) (FWE corrected *P*s < 0.001 one-sided), which correspond to the blue and yellow areas in Figure 3A.

Since, the utility representation for use of interpersonal comparison must represent utility irrespective of varying reward type, we sought to identify the brain region that represents the utility prediction errors for both food and money. Most of the ACC/vmPFC region that represents the utility prediction error for food also correlated with the utility prediction error for money, hereafter called the “utility region” (the yellow area in Figure 3A). Thus, the “utility region” correlated with the prediction error of utility rather than that of the reward amount irrespective of reward type. In this paper, we refer to the PSCs of the utility region as Δu_neural_.

### Interpersonal comparison of utility based on neural signals

To realize an interpersonal comparison of utility based on each person’s household income, we needed to determine the relationship between income and Δu_neural_. We applied a multiple regression analysis with regressors for income, age, and sex. The results showed that Δu_neural_ was significantly affected by income (β_income-1_ = 0.247 ± 0.132 (mean ± SE), t(56) = 1.874, *P* = 0.033 one-sided; Figure 3C) but not the other regressors (β_age_ = -0.100 ± 0.132, t(56) = -0.757, *P* = 0.452 two-sided; β_sex_ = -0.095 ± 0.130, t(56) = -0.731, *P* = 0.468 two-sided; β_0_ = -0.235 ± 0.179, t(56) = -1.318, *P* = 0.193 two-sided). This indicated that Δu_neural_ was inversely proportional to income, supporting the hypothesis that the larger the income, the smaller the Δu_neural_.

In the above analysis, we assumed that utility of income can be approximated by a logarithmic function (i.e., Δu_neural_ was assumed to be inversely proportional to income). Since this assumption was arbitrary, we compared Δu_neural_ between the participants with a household income up to 6 million yen (the “poor” group) and those with a household income above 6 million yen (the “rich” group) (Figure 3B) to subsequently consider a distribution method for the MRI participants that did not require any specific utility function (see *Application to a distribution problem*). We found a significant difference in Δu_neural_ between the two income groups (t(58) = 2.39, *P* = 0.010 one-sided, BF_+0_ = 5.723, median posterior δ = 0.535 with 95% CI [0.046, 1.046]) (Figure 3D) but no difference in subjective probabilities (Supplementary Figure 4). The posterior means of Δu_neural_ across participants, E[μ_PSCpoor_] and E[μ_PSCrich_], were 4.624 * 10^-4^ and 2.007 * 10^-4^, respectively (E[μ_PSCpoor_]/E[μ_PSCrich_] was 2.304). Thus, we were able to objectively determine an *inter*personally comparable scale of utility difference by using Δu_neural_ (Figure 1D). The scale of utility difference would not have been interpersonally comparable if it was based on choice behavior alone.

### Coincidence of the ratio of Δu_neural_ and impartial spectators’ estimation

To validate our scientific interpersonal comparison method using Δu_neural_, we developed a behavioral task to measure impartial spectators’ estimates of the utility difference ratio of the poor and the rich group. An additional 15 participants were recruited, from the university we used to recruit participants for the gambling experiment, completed the impartial spectator task. Two participants were excluded from the analysis, because they did not pass the instructional manipulation check (see *Methods*). We considered the remaining 13 participants “impartial spectators” for this task, because they were impartial and empathetic to the MRI participants (Harsanyi 1977; Smith 1759) (see *Methods*). The impartial spectator participants were presented with a histogram showing the household income distribution of the MRI participants in the gambling experiment and were asked to give their intuitive estimates of the amount of money needed for participants in the rich group to experience the same level of pleasure as those in the poor group would when they received ¥400, ¥500, or ¥600 (see *Methods*). The responses are shown in Figure 4A. The impartial spectator participants’ intuitive estimates of the utility difference ratio of the poor and the rich group were calculated (median, 2.000) (Figure 4B). The utility difference ratio of the two groups based on the impartial spectator participants’ intuitive estimates coincided with those based on Δu_neural_ (E[μ_PSCpoor_]/E[μ_PSCrich_] = 2.304) with moderate evidence (BF_10_ = 0.165, median posterior δ = 0.014 with 95% CI [-0.456, 0.211]) (Figures 4B; Supplementary Figure 5). Thus, the interpersonal comparison method based on MRI signals from the brain was coherent to the impartial spectators’ estimation. This result means that our new method for interpersonal comparison of utility is consistent with the conventional intuitive method, and is also a superior method that is objective and reproducible.

**Figure 4.**
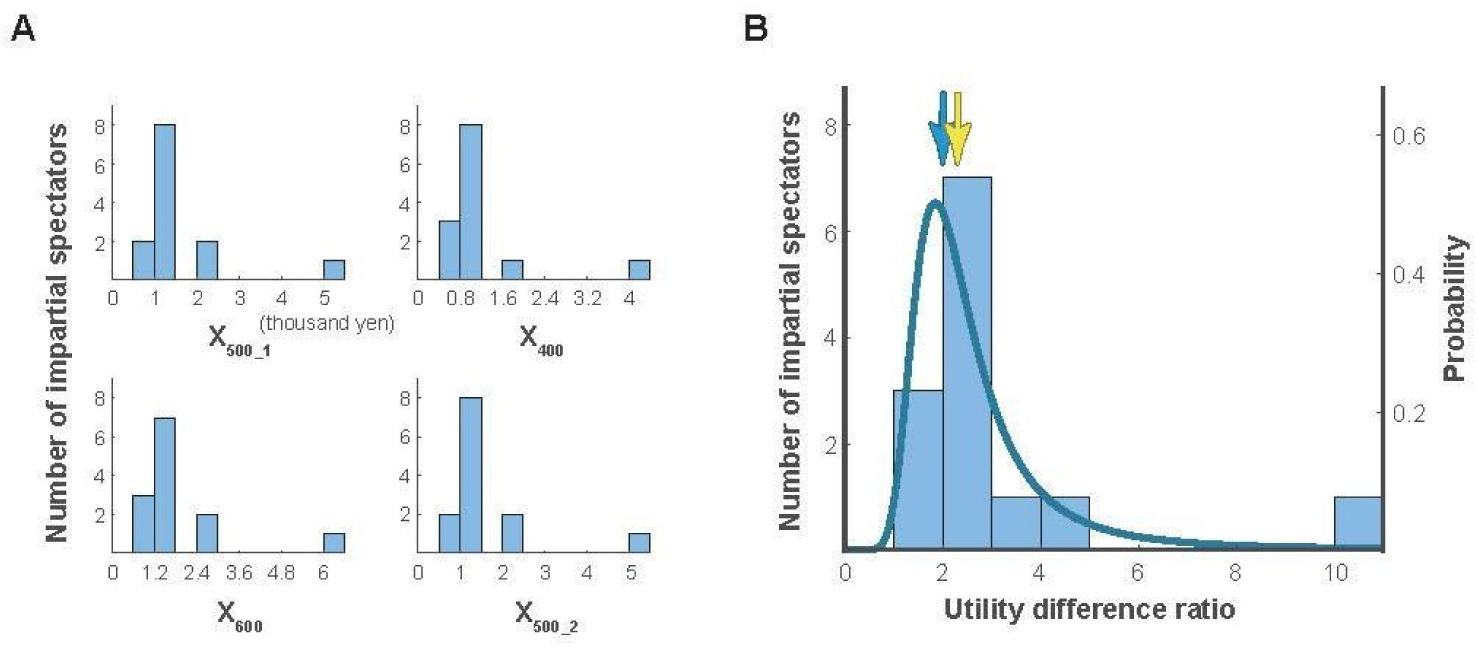
Validation by impartial spectators’ estimation. (**A**) The distribution of impartial spectators’ estimates of the equivalent amounts of money needed to please the rich group based on the poor group of participants’ receipt of ¥500, ¥400, ¥600, and ¥500 in the impartial spectator task. X_500_1_, X_400_, X_600_, and X_500_2_ are the estimates for the corresponding questions. (**B**) The distribution of the impartial spectators’ estimates of the utility difference ratio of the poor group to the rich group based on the combined data of **A** and its fit curve on the Fieller-Hinkley distribution. Yellow and light blue arrows indicate the Δu_neural_ ratio and the median of impartial spectators’ estimates, respectively.

### External validity check using the ABCD study’s data

To validate our method in a larger independent sample, we analyzed the data of 5,372 children (aged 9–10 years, living in the USA) who had participated in the ABCD study (Casey et al. 2018). Since we had already established that Δu_neural_ calculated in the gambling task was useful for an interpersonal comparison, we analyzed the data obtained in the monetary incentive delay (MID) task (Knutson et al. 2001) in which Δu_neural_ was available as well. We focused on Δu_neural_ (here PSC in the utility region per 1 dollar) during the feedback phase of the MID task. We employed a linear mixed model with the study site as a random effect to account for potential site-specific variations. The model included the reciprocal of household income, sex, and age as fixed effects. We found a significant positive association between the reciprocal of household income and neural activity in the utility region (β_income-1_ = 0.054 ± 0.014, z = 3.939, *P* < 0.001 two-sided; Figure 5). This result aligned with our main findings, supporting the inverse relationship between income and Δu_neural_. Additionally, we observed significant effects for sex (β = -0.092 ± 0.027, z = - 3.405, *P* = 0.001 two-sided, for female participants compared to male participants) and age (β = -0.031 ± 0.014, z = -2.301, *P* = 0.021 two-sided), indicating that these factors also influenced utility-related neural responses in children.

**Figure 5.**
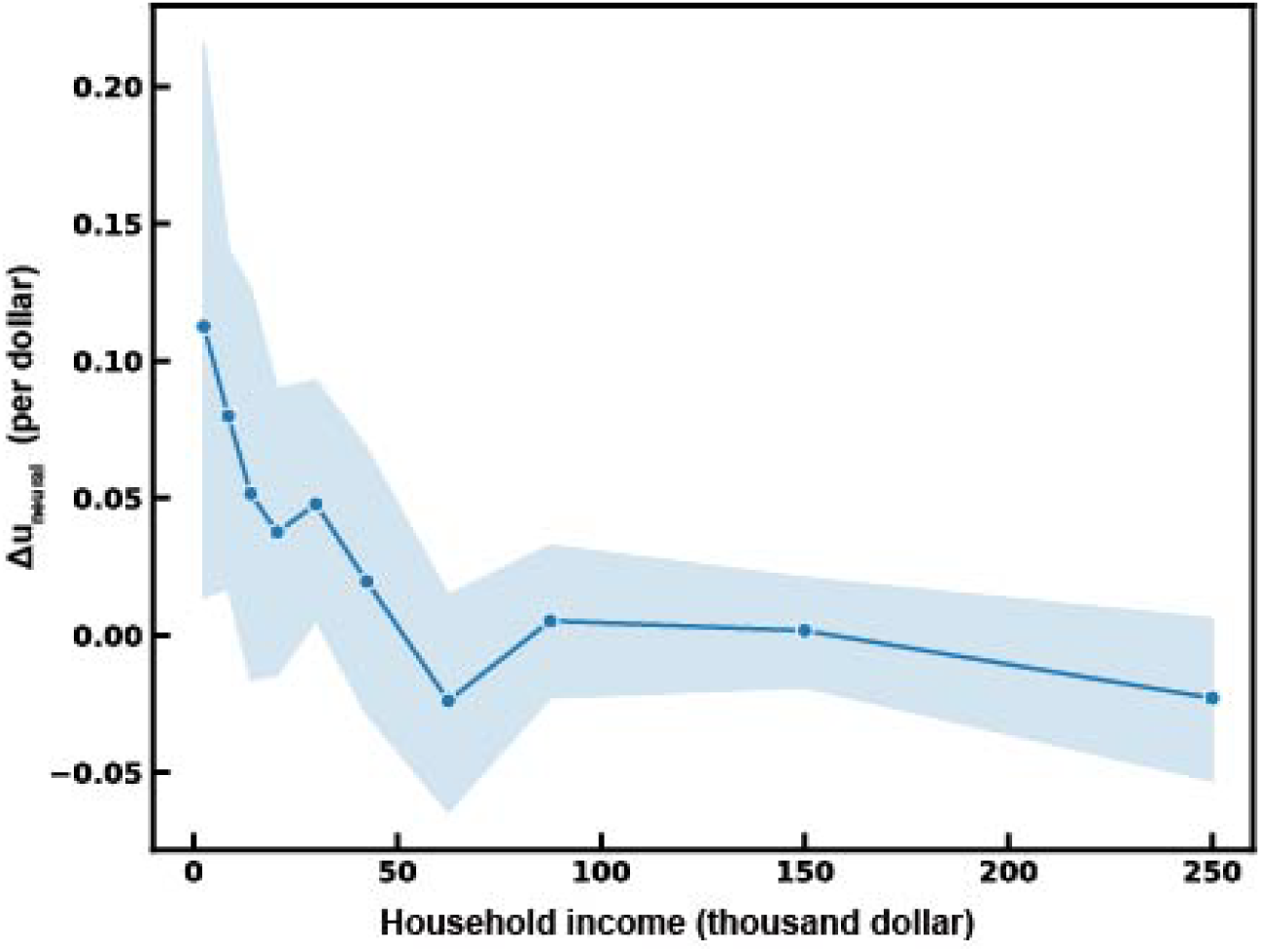
External validity check of interpersonal comparisons with large scale fMRI study. The Δu_neural_ of participants plotted against their household income for 5,372 children from the ABCD study. Δu_neural_ represents the PSC per $1 in the utility region during the feedback phase of the MID task. Δu_neural_ was inversely proportional to income (βincome^-1^ = 0.054 ± 0.014 (mean ± SE), z = 3.939, *P* < 0.001 two-sided) after regressing out age, sex, and study site. The shaded area indicates the 95% confidence interval of the regression line.

To characterize the population-level utility function with respect to income, as measured by Δu_neural_, we employed a nonlinear model fit approach using an isoelastic utility function. This method yielded an estimated elasticity parameter of 0.662 (95% CI: 0.004 - 1.137). This value provided a concise summary of the curvature of the utility function in the studied population.

The results of the ABCD study provided external validation for our main findings, demonstrating that the inverse relationship between income and utility-related brain activity is observable in a large, diverse sample of children aged 9–10 years living in the USA. This extends our understanding by showing that this neurally decoded utility is present from an early age in a different culture, suggesting a universal and early emerging aspect of how the brain represents utility across socioeconomic strata.

### Application to a distribution problem

Finally, we applied the interpersonal comparison method based on Δu_neural_ to a distribution problem from a social planner’s perspective (right balloon in Figure 6) for the participants in the gambling experiment. Here, we used a decision rule that maximized utilitarian social welfare, because a utilitarian social welfare function is most common when utility differences are interpersonally comparable (d’Aspremont and Gevers 1977; Narens and Skyrms 2020; Roberts 1980). Suppose a case in which a planner distributes ¥1,000 to all participants (poor: n = 30, rich: n = 30) in addition to their usual income (Policy A). Since E[μ_PSCpoor_] and E[μ_PSCrich_] are 4.624 * 10^-4^ and 2.007 * 10^-4^, respectively, the expected increase in social welfare would be 19.893 util (= (4.624 + 2.007) * 10^-4^ * 1,000 * 30), assuming that 1 util corresponds to a 1% change in the BOLD signal. Next, we considered a case in which the planner distributes a certain amount of money selectively to the 30 poor participants. For the selection process, we included a regulation cost. Suppose a case in which the planner distributes ¥1,500 to every poor participant with the assumption of a regulation cost of ¥500 (Policy B). In this case, the expected social welfare increase will be 20.808 util (= 4.624 * 10^-4^ * 1,500 * 30) for Policy B. Because the social welfare increase is higher for Policy B than Policy A, we decided to use Policy B to distribute ¥1,500 to every poor participant instead of ¥1,000 to all participants, and actually carried this out for the participants who agreed to receive additional compensation. Without an interpersonal comparison of utility, we cannot argue that such a greater increase in social welfare will result from a policy that is more generous to the poor than to the rich.

**Figure 6.**
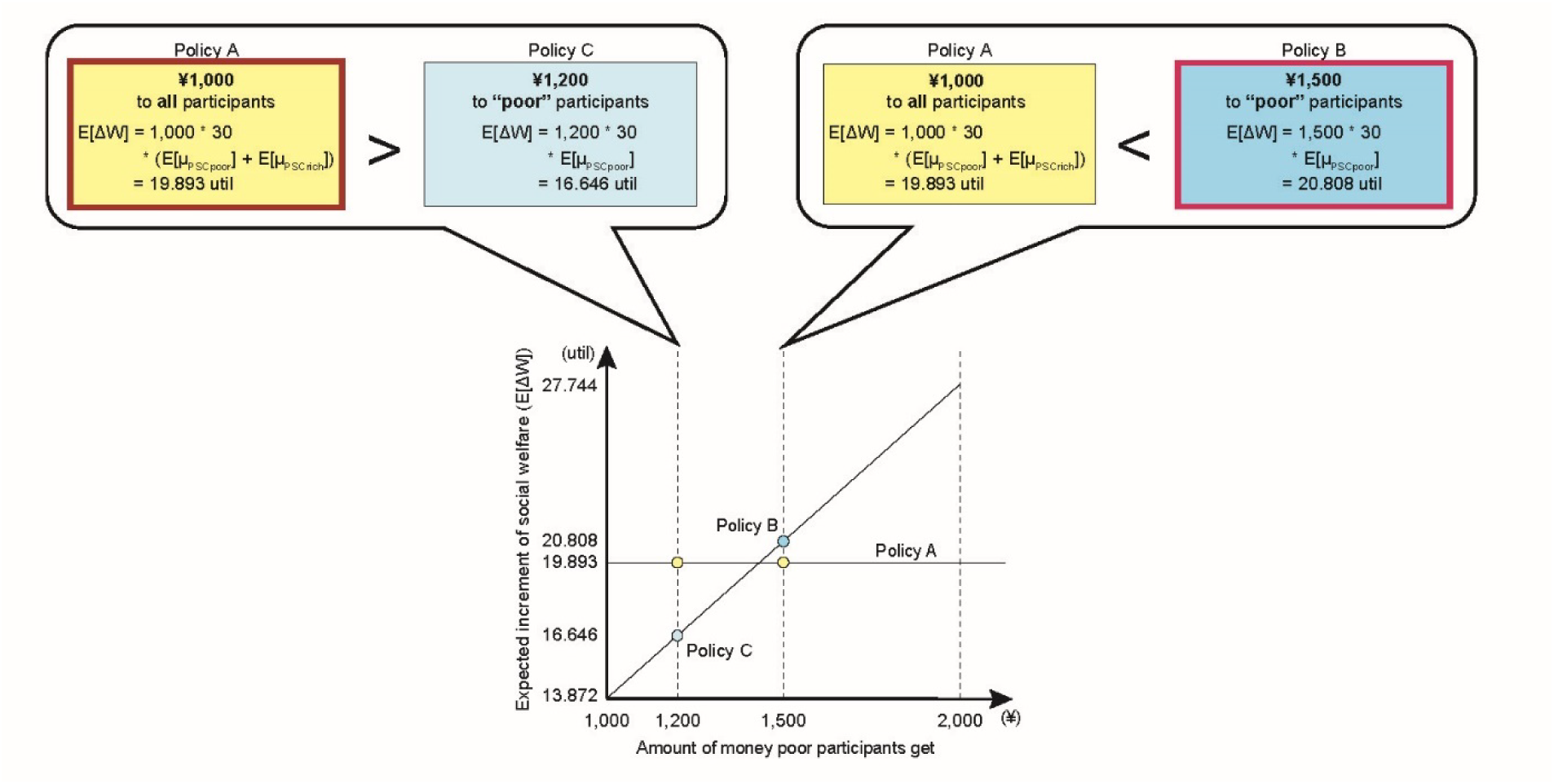
Application to a distribution problem. The two distribution policies, A and B, are compared (right balloon). Policy B was chosen, because it increased expected utilitarian social welfare (E[ΔW]) more than Policy A. Policies A and C were also compared (left balloon). Policy A would be chosen over Policy C using the same rationale. The plot illustrates expected increments of social welfare by distributing ¥1,000 to all participants (poor: n = 30, rich: n = 30) and by distributing ¥x to “poor” participants, including points for the compared policies A, B, and C.

According to the ratio of posterior means of Δu_neural_ (E[μ_PSCpoor_]/E[μ_PSCrich_] = 2.304) described above, for the problem of choosing between a policy that distributes ¥k to all the participants and a policy that distributes ¥(k + *ρ*) to every poor group participant, the optimal decision rule for an expected utilitarian social welfare maximizer can be generalized as follows:

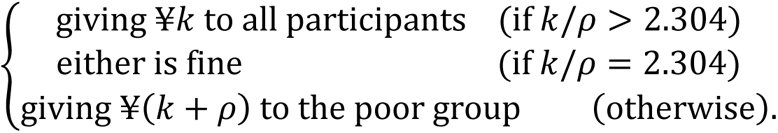

This decision rule tells us that the increase in social welfare that results from distributing ¥1,200 to every poor participant with the assumption of a regulation cost of ¥800, would be less than giving ¥1,000 to all the participants (left balloon in Figure 6). Thus, by finding a scientific method for interpersonal comparisons of utility, we have succeeded in overcoming Arrow’s impossibility theorem and solving a distribution problem.

## Discussion

To develop a scientific method for an interpersonal comparison of utility, we measured the functional MR signals of participants with different household incomes while they were performing food and monetary gambling tasks. We obtained the following results. First, activities in an ACC/vmPFC region were correlated with the prediction error of utility irrespective of reward type. Second, the signal strength from this region of the brain (Δu_neural_) was inversely proportional to household income across all the participants. Third, the ratio of Δu_neural_ in lower- and higher-income participants coincided with estimates of their psychological pleasure by impartial spectators. Fourth, we validated our findings using data from over 5,000 children in the ABCD study, confirming the inverse relationship between Δu_neural_ and household income in a large, diverse sample. Finally, on the basis of the ratio of Δu_neural_, we actually solved a distribution problem.

We succeeded in developing a scientific method for interpersonal comparisons of utility based on activities in a specific region of the brain (ACC/vmPFC). Scientific measurement methods are necessary for interpersonal comparisons of utility, because interpersonal comparison methods based solely on impartial spectators’ estimates are unlikely to work well when judgments are polluted by systematic biases (Chang 2004; Müller-Lyer 1889). Without scientific measurement methods, we cannot determine whether systematic biases are present. Furthermore, non-scientific interpersonal comparison methods based on empathy may work in a small group but not in society as a whole, because empathy requires too rich information to reach people living in different cultural contexts (Bloom 2016; Elster and Roemer 1991). Our new interpersonal comparison method based on a scientific measurement (fMRI) extends empathy and be applicable to society as a whole, as demonstrated across diverse samples in the large-scale ABCD study.

Our finding that the activity of the ACC/vmPFC correlated with the prediction error of utility rather than that of the objective amount of reward is consistent with previous studies that indicate the ACC/vmPFC encodes subjective value (Bartra et al. 2013; Glimcher and Fehr 2013; Matsumoto et al. 2007); however, in contrast to these previous studies, the striatum was not extracted in our study. This may be due to differences in the respective methodologies. We extracted brain activity that correlated with utility during the food gambling task in particular by using a rigorous method to determine the utility curve (i.e., the fractile method); therefore, it is possible that the ACC/vmPFC specifically represents the prediction error of utility but the striatum does not. This interpretation is consistent with our finding that striatum activation was superficially associated with the prediction error of utility in the monetary gambling task (Figure 3A); this is likely due to the linearity of the utility function of money, which prevented us from distinguishing between the prediction error of utility and that of the reward amount represented in the striatum (Seymour et al. 2007). Previous studies have also suggested that the ACC/vmPFC represents more subjective information than the striatum (Aoki et al. 2014; Murayama et al. 2015).

A previous study showed that midbrain dopamine neurons encode the prediction error of utility, not the reward amount, using the fractile method in monkeys (Stauffer et al. 2014). Dopaminergic inputs may supply information about prediction error of utility in ACC/vmPFC in humans, because the ACC/vmPFC receives inputs from dopamine neurons in both monkeys and humans (Gaspar et al. 1989; Lewis et al. 1988).

We identified the “utility region” in the ACC/vmPFC by extracting the brain region that represented utility prediction errors in both food and monetary gambling tasks. This is consistent with previous neuroimaging studies that reported that specific brain regions, including the ACC/vmPFC, represent the value of various rewards (e.g., food, money, and social praise) on a single common scale (Bartra et al. 2013; Glimcher and Fehr 2013; Izuma et al. 2008; Levy and Glimcher 2012) like utility in economics.

One study argued that inter-individual (or inter-group) differences in fMRI studies are difficult to interpret if the variances of the explanatory variables differ between groups (Lebreton et al. 2019); however, that was not the case in the present study. In the monetary gambling task, the variances of the explanatory variable (i.e., prediction error of monetary amount) did not different between the poor and the rich group, because all offers were common among the participants, and the subjective probabilities did not differ between the poor and the rich group (Supplementary Figure 4).

Our interpersonal comparison method can be applied to other kinds of goods by following certain procedures, although we used only monetary and food rewards in our experiment. First, willingness to pay for other goods can be measured to find out how much the good is worth to the person, and then willingness to pay can be converted to the interpersonally comparable utility of other goods by multiplying the coefficients of the utility measured using our proposed method.

We did not find a significant modulation of ACC/vmPFC activity by the two reward range (narrow and wide) conditions in the monetary gambling task. This result indicated that the ACC/vmPFC activity was robust to different ranges of reward amounts in our experiment. However, we do not argue that this activity is totally free from the normalization that previous studies have suggested (Burke et al. 2016; Kobayashi et al. 2010; Louie et al. 2015). The apparent lack of influence by changes between conditions might be due to the present study’s experimental design. The task consisted of three sessions of 40 trials (20 successive trials each for the narrow and the wide condition) with a 1 min break between the sessions. We explicitly informed the participants of this when presenting the instructions for the monetary gambling task, which may have caused the participants to treat the narrow–wide structured trials as a single block of trials and suppressed the influence of changes between the conditions. The univariate method we applied to analyze the brain data may also have caused robustness to the changes between conditions (Burke et al. 2016). A multivariate analysis may be more efficient for detecting signals that are sensitive to such changes (Burke et al. 2016; Woo et al. 2017).

In the present study, we found a neural representation of utility prediction error. Some readers may think that neural activity in the utility region is also a function on the consequential states, because utility functions are usually defined on consequential states and not on the processes that lead to them. However, previous studies have shown that the activity of the reward system, including vmPFC, is modulated not only by consequential states (Bartra et al. 2013) but also by processes like opportunity equality (Aoki et al. 2014). Thus, it is unlikely that utility-related information in the brain is limited to the consequential states usually considered in utility functions.

In addition to our primary experiment, we validated our findings using a larger independent sample from the ABCD study, which included over 5,000 children. The analysis focused on Δu_neural_ during the feedback phase of the MID task. We observed that Δu_neural_ was inversely related to household income, aligning with the results of our primary analysis. This consistency across different age groups and cultures using a much larger sample size strongly supports the robustness of our method for interpersonal comparisons of utility based on brain activity. The fact that we detected similar patterns in children from diverse socioeconomic backgrounds suggests that the neural basis of utility representation and its relationship with income are established early and remain relatively stable (Rakesh et al. 2023). This broad external validation suggests that our method for interpersonal comparisons of utility based on brain activity is applicable across different demographic groups and potentially offers new insights into socioeconomic influences on neural processing. Furthermore, the universal neural representation of utility, formed from early developmental stages, offers valuable implications for education and welfare.

In the present study, we applied univariate analyses to read utility signals, because they are not only more straightforward than multivariate analyses but also seemed more efficient for detecting signals that are not normalized across contexts (Burke et al. 2016); however, there may be a multivariate analysis that improves the accuracy of reading out utility signals (Woo et al. 2017).

Another justification for interpersonally comparable utility based on neural activity could be its ability to genuinely explain something additional, such as fitness. The neural activity-dependent utility may appear through a variety of outputs such as facial expression and subtle movements as well as choice behaviors. These outputs may be detected by other consanguineous individuals and evoke their caring behaviors that affect fitness. By providing a scientific basis for comparing utility across individuals, our study opens up new avenues for exploring the relationship between neural representations of utility and evolutionary fitness. While utility has generally been considered incomparable between individuals, fitness is viewed as interpersonally comparable. Our method for an interpersonal comparison of utility based on neural activity potentially bridges this gap between utility and fitness (Fehr and Camerer 2007). Thus, this approach could reveal both alignments and divergences between individual utility and adaptive outcomes, thereby offering insights into their complex relationship at a population level (Okaha et al., 2014) and facilitating the integration of evolutionary theory and decision theory. However, it should be noted that in the highly cultural environment in which modern humans live, the link between individual utility and fitness has weakened (Sterelny 2012); consequently, human individuals may pursue actions that do not contribute to their fitness.

In the current study, we focused on utility difference, not on utility level, to compare utility interpersonally, because it could be treated quantitatively in the experiment. By comparing utility differences interpersonally, we conducted a redistribution to maximize the utilitarian social welfare of the participants within our budget constraints. Here, we note that the present study itself does not derive a normative conclusion on distributive justice (Hume’s law, Hume 1739). Utility level is also important to realize a full version of interpersonal comparison of utility (d’Aspremont and Gevers 1977; Roberts 1980; Sen 2018). Future studies should conduct interpersonal comparisons of utility levels, which enables distributions according to the variable principles discussed in modern justice theories other than utilitarianism (d’Aspremont and Gevers 1977; Hirose 2014; Roberts 1980).

The process of constructing a quantitative concept and standardizing its measurement methods should be viewed as progressive processes that increase coherence among the elements of theory and instrumentation (Chang 2004; Tal 2013, 2015). This is also the case when constructing an interpersonally comparable utility. The problem of measuring utility has frequently been compared with the problem of measuring temperature (Arrow 1963; Von Neumann and Morgenstern 2007). We validated our interpersonal comparison method by showing coherence between objective measures (i.e., fMRI signals) and human intuition (Goldman 1995; Harsanyi 1977; Rossi 2014), as was done for the temperature scale in its initial stages (Chang 2004).

The proposed method is significant in an academic sense, since it enables us to escape from Arrow’s impossibility (Arrow 1963; Sen 2018); moreover, it can be applied for evidence-based policy-making in nations that use cost-benefit analyses or optimal taxation theory for policy evaluation (Boardman et al. 2018; Saez 2001). In this study, we demonstrated a major advance in social science using a neuroscientific measure (Camerer et al. 2005). Although the present study does not draw normative conclusions (Hume’s law) on distributive justice, it does constitute a significant epistemological advance as a starting point for subsequent scientific inquiries (Chang 2004) to standardize a measurement method for utility as an interpersonally comparable quantity and for philosophical reflections on social structures (Daniels 1979; Rawls 1971).

## Methods

### Gambling experiment: participants

Sixty-three healthy right-handed student volunteers who live with their parents participated in this experiment. All the participants were native Japanese speakers at Tamagawa University and provided their written informed consent prior to participation. None of the participant had a history of neurological or psychiatric disease. The study was approved by the ethics committee of Tamagawa University in accordance with the ethical standards laid down in the 1964 Declaration of Helsinki and its later amendments. One participant was excluded from the analysis because of a technical problem with data storage. In the monetary gambling task, two participants were excluded from the analysis because of excessive head motion (> 3 mm) during the task, resulting in the final sample of 60 (35 female and 25 male participants; age: 20.13 ± 0.98 years; range: 18–22 years). In the food gambling task, three participants (two were the same participants excluded from the monetary gambling task) were excluded because of excessive head motion, and three more participants were excluded because of insufficient motivation to obtain the food tickets, based on < 5 certainty equivalents (CEs) at largest. This left a final sample of 56 participants (33 female and 23 male participants; age: 20.09 ± 0.98 years; range: 18–22 years).

### Procedures

The participants completed monetary and food gambling tasks in an MRI scanner. Linear utility functions were assumed for money, and non-linear utility functions were assumed for food. Afterward, participants completed a probability weighting task outside the MRI scanner to measure subjective probability for a 50/50 gambling situation.

### Monetary gambling task

In this task, participants had to choose between a sure (riskless) offer and a lottery (Figure 2C). The participants were told they could gain or lose money in each trial depending on the outcome of the gamble. Each fMRI session consisted of a narrow block (20 trials) in which the money outcomes were in the range ¥-200∼200 and a wide block (20 trials) in which the money outcomes were in the range ¥-400∼400. Participants were explicitly informed of each block’s range of possible monetary outcomes. The current block was identified by a color disk on which the amount of money was displayed.

Each trial began with a fixation cross at the center of the monitor (2 s) followed by a sure offer (2 s). There were five sure offers in the narrow block (¥-100,-50, 0, 50, and 100) and five in the wide block (¥-200,-100, 0, 100, and 200). Next, the sure offer disappeared and the lottery offer was presented (2 s). Each lottery offer was a gamble entailing a 50% chance of gaining (or losing) an amount of money and a 50% chance of neither gaining nor losing the money (¥0). There were four lottery amounts in the narrow block (¥-200, -100, 100, 200) and four in the wide block (¥-400, -200, 200, 400). Next, the lottery offer disappeared and participants chose between the sure and lottery offers. Participants had to press a left or right button with the index or middle finger of their right hand within 30 s to choose the corresponding offer. The trial was aborted if the participant failed to respond within the time window (“error trial”). The association between the buttons and offers was counterbalanced across participants. Chosen offers were presented on the monitor immediately after pressing the button. Next, the amount of money earned was displayed in flashing yellow text.

The delay between selection (button press) and feedback (offer) was jittered between 3 and 7 s (average 5 s). Inter-trial intervals were jittered between 3 and 7 s (average 5 s). All 5 * 4 = 20 possible combinations of sure and lottery offers were presented in randomized order in each block. Each participant was engaged in this task for 3 sessions of 40 trials except the error trials. The order and color of the two blocks were counterbalanced across participants. The block-start cue (“Next block is narrow” or “Next block is wide”) was presented at the beginning of each block (5 s). Stimulus presentation and timing data collection for all stimuli and response events were performed using MATLAB (MathWorks, Natick, MA, USA) and Psychtoolbox (www.psychtoolbox.org).

Previous studies have shown that the utility function for money is approximately linear for moderate amounts of money (Abdellaoui et al. 2008; Booij and Kuilen 2009; Wakker and Deneffe 1996). We regarded the utility function for money as linear, because we used small amounts of money. We used a food gambling task to dissociate the brain regions that represents utility from the amount of the reward.

### Food gambling task

In the food gambling task, each participant had to choose between a sure (riskless) offer and a lottery with the same appearance as the monetary gambling task (Figure 2A). In this task, participants could receive tickets that could be exchanged for snacks. Before the fMRI scan, each participant chose their preferred snack (rice cracker or chocolate). Participants were instructed that at the end of the experiment, one trial would be randomly selected and the number of tickets that could be exchanged for their preferred snack would be determined according to the outcome of their actual decision on that trial. One ticket could be exchanged for one snack, so the number of tickets they obtained was the maximum number of their preferred snack they could receive.

Each trial began with a fixation cross at the center of the monitor (2 s) followed by the sure offer (2 s). The possible number of tickets in the sure offers was 1∼300. Then the lottery offer was presented (2 s). Each lottery offer was a gamble entailing a 50% chance of gaining one of two numbers of tickets. The possible number of tickets in lottery offers was 1∼300. Then, the lottery offer disappeared and participants chose between the sure and lottery offers. Each participant was required to press the left or right button with the index or middle finger of their right hand within 30 s to choose the corresponding offer. The trial was aborted if the participant failed to respond within the time window (“error trial”). The association between the buttons and offers was the same as in the monetary gambling task. The chosen offer was presented on the monitor immediately after pressing the button. Then, the numerical outcome from the chosen offer was displayed in flashing yellow text. The delay between selection (button press) and feedback (offer) was jittered between 3 and 7 s (average 5 s). Inter-trial intervals were jittered between 3 and 7 s (average 5 s). The color of the disk (red or blue) was counterbalanced across participants. In this task, we measured certainty equivalents (CEs) based on the fractile method with three phases (Supplementary Figure 2B) (see *Constructing utility functions in the food gambling task*). In each phase, the CE measurement trials and random trial (all three numbers were sampled from a uniform distribution, unif{1, 300}) were randomly permuted and this procedure was repeated until all CEs converged (see *Behavior analyses: measuring CEs*). Each participant was engaged in this task for 3∼5 sessions of 40 trials and completed a minimum of 120 trials. The fMRI data from these 120 trials were analyzed, whereas all the trials were used to analyze the behavioral data. The fMRI session for this task was terminated when the sum of the completed trials from the food task and the error trials from both monetary and food tasks reached 200. Stimulus presentation and timing data collection for all the stimuli and response events were performed using MATLAB (MathWorks, Natick, MA, USA) and the Psychtoolbox (www.psychtoolbox.org).

### Probability weighting task

This task was very similar to the previous tasks. Participants were instructed that at the end of the experiment, one trial would be randomly selected, and a monetary payment would be made according to their decision. We measured the CE of the 50/50 gamble (*p* = 0.5 for each outcome) between ¥2,000 and ¥2,500 using parameter estimation through sequential testing (PEST) (see *Behavior analyses: measuring CEs*). The CE measurement trials and the random trial (all three numbers were sampled from a uniform distribution, 10 × unif{200, 250}) were randomly permuted. This procedure was repeated until the CE converged. Each participant engaged in a task for typically 30–40 trials. The stimulus presentation and timing data collection for all the stimuli and response events were performed using MATLAB (MathWorks, Natick, MA, USA) and Psychtoolbox (www.psychtoolbox.org).

### Questionnaire

After the gambling experiment, each participant was asked about their understanding of the tasks and household income.

### MRI data acquisition

Functional imaging was conducted using a 3T Trio A Tim MRI scanner (Siemens, Germany) and gradient echo T2*-weighted echo-planar images (EPI) were acquired with blood oxygenation level-dependent contrasts. Forty-two contiguously interleaved transverse EPI slices were acquired for each volume (slice thickness, 3 mm; no gap; repetition time, 2,500 ms; echo time, 25 ms; flip angle, 90°; field of view, 192 mm^2^; matrix, 64 × 64). Slice orientation was tilted −30° from the AC-PC line. For each participant, data were acquired in three scanning sessions for the monetary gambling task and 3–5 scanning sessions for the food gambling task. We discarded the initial three dummy volumes per session without analysis to allow for the T1 equilibrium effect. High-resolution anatomical T1-weighted images were acquired for each participant.

### Behavioral analyses: measuring CEs

We used PEST to measure CEs in food gambling and probability weighting tasks (Luce 2000; Stauffer et al. 2014) (Supplementary Figure 2A). We assessed the number of tickets or the amount of money in the sure offer that was subjectively equivalent to the value associated with each lottery offer. The rules governing the PEST procedure were adapted from Luce (2000). Each trial consisted of three numbers: a sure number (SU), a lottery win number (LW), and a lottery loss number (LL). Each PEST sequence consisted of several trials during which one constant lottery offer (constant LW and LL) was presented as a choice option against the sure offer (variable SU).

In the initial trial of the PEST sequence, SU was determined by rounding (LW + LL)/2 to the nearest integer. Based on the participant’s choice between the sure offer and the lottery, SU was adjusted in the subsequent trial. If the participant chose the lottery offer in trial *t*, then SU was increased by ε in trial *t* + 1; however, if the participant chose the sure offer in trial *t*, SU was reduced by ε in trial *t* + 1. Initially, ε was determined by rounding (LW + LL)/4 to the nearest integer and adjusted according to the following rules. Rule 1: After the third trial of a PEST sequence, every time the participant switched from one option to the other, the size of ε was halved. Rule 2: If the updated SU based on ε was equal to LW or LL, the size of ε was halved and then SU was updated. Thus, the procedure converged by locating subsequent sure offers on either side of the true indifference value and reducing ε until the interval containing the indifference value was small (1 for tickets and 10 for money). When the updated SU was the same as the previous SU, the PEST procedure terminated.

### Constructing utility functions in the food gambling task

We determined each participant’s utility function for food tickets in a range between 1 and 300 tickets using the fractile method based on 50/50 gambles (Stauffer et al. 2014); see Supplementary Figure 2B for an example the fractile procedure. To do so, we first measured the CE of a 50/50 gamble between 1 and 300 tickets (x_1_) using PEST. We then used x_1_ as an outcome to construct two new gambles (1 ticket, *p* = 0.5 and x_1_ ticket, *p* = 0.5; x_1_ ticket, *p* = 0.5 and 300 tickets, *p* = 0.5) and measured their CEs (x_2_ and x_3_). Finally, we measured the CEs of a 50/50 gamble between 1 and x_2_ tickets (x_4_), x_2_ and x_1_ tickets (x_5_), x_1_ and x_3_ tickets (x_6_), and x_3_ and 300 tickets (x_7_). We fitted the CE data using local data interpolation (i.e., splines, MATLAB SLM tool) (Stauffer et al. 2014), after determining the probability weighting function. This procedure fitted cubic functions on consecutive segments of the data.

We used two polynomial pieces (three knots) for our fittings. The left knot was located at 1, the right knot was located at 300, and the middle knot’s position was determined by minimizing the difference between the empirical data and the fitted curve (the candidate positions were 2, 3, … 299). We restricted the fitting to have a positive slope over the entire range of outcomes. We required the function to be increasing weakly based on the fundamental economic assumption that more goods do not decrease total utility. This procedure provided an estimation of the shape of the participant’s utility function in the range of 1–300 tickets.

### Estimating w(*p* = 0.5) in probability weighting task

We set u(¥x) = αx + β based on the assumption that utility for moderate amounts of money is approximately linear. This means that the value of the probability weighting function *w* when *p* = 0.5 can be measured using the CE in the probability weighting task, according to the following equations:

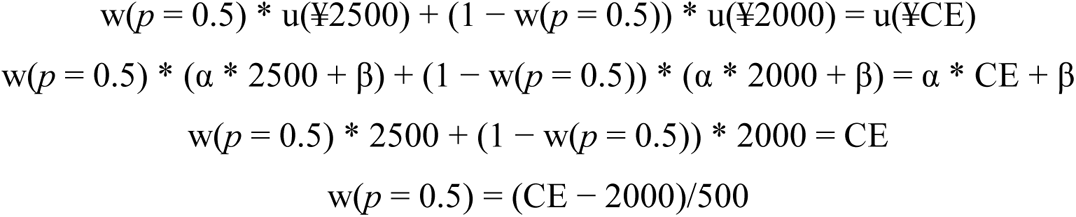

Note that these equations hold in both original prospect theory (Kahneman and Tversky 1979) and cumulative prospect theory (Tversky and Kahneman 1992), because the two theories coincide in the domain of two-outcome lotteries (Abdellaoui et al. 2007). The expected utility of a lottery with probability weighting function can be calculated as follows:

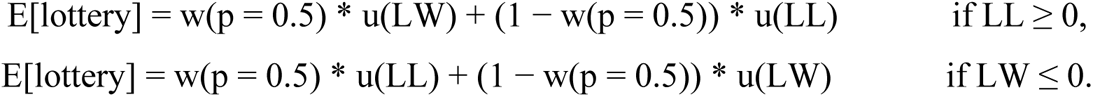

We compared the values of w(*p* = 0.5) of the participants whose household income placed them in the “poor” group and those in the “rich” group using the classical t-test and Bayesian t-test with default effect size priors (Cauchy scale 2^0.5^/2) (Keysers et al. 2020) (Supplementary Figure 4). BFs were computed using the Savage–Dickey density ratio method. Note that BF < 1/3 is considered moderate evidence for the null hypothesis and BF > 3 is considered moderate evidence for the alternative hypothesis (roughly similar to *p* < 0.05) (Keysers et al. 2020; Lee and Wagenmakers 2013). Bayesian t-test results were reported using the two-tailed Bayes factor BF_10_ that represents P(data| H_1_: μ_wpoor_ ≠ μ_wrich_)/P(data| H_0_: μ_wpoor_ = μ_wrich_), where μ_wpoor_ and μ_wrich_ indicate the mean of w(*p* = 0.5) in the “poor” and the “rich” group, respectively. Effect size estimates were reported as median posterior Cohen’s δ with 95% CI. Bayesian hypothesis tests were performed using R (https://www.R-project.org/) and Stan (https://mc-stan.org/), and all Markov chain Monte Carlo settings were 3 chains with 100,000 samples each, with a burn-in of 10,000 and a thinning factor of 1. For all parameters, 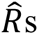 were lower than 1.1.

### Validation of participants’ utility functions

We validated the shape of each participant’s utility function (estimated curve for food tickets) in the food task using random trials in which the choice options did not depend on the participants’ previous choices, that is, incentive compatible (Stauffer et al. 2014). In these trials, we assessed the percentage of trials in which participants chose the offer with a larger estimated utility and compared them with those assessed based on the expected reward amount weighted by subjective probability using the classical t-test.

### Preprocessing of functional images

The following preprocessing procedures were performed using statistical parametric mapping (SPM12) software (Wellcome Department of Imaging Neuroscience, London, UK; https://www.fil.ion.ucl.ac.uk/spm/) implemented in MATLAB with correction for head motion, adjustment of acquisition timing across slices, spatial normalization to the standard MNI template, and smoothing using a Gaussian kernel with a full-width at half maximum of 8 mm.

### Functional MRI data analysis

We adopted a conventional two-level approach using SPM12 to analyze the fMRI data. A set of regressors was generated by convolving a canonical hemodynamic response function provided by SPM12 with a series of epochs (initial fixation, sure offer, lottery offer, choice, and feedback). We applied a voxel-by-voxel multiple regression analysis to the preprocessed images for each participant. Statistical inference on contrasts of the parameter estimates was then performed at the second-level between-participant analysis, using a one-sample t-test. As we had an a priori hypothesis focusing on the activations of the value system, the second-level analysis was performed for the voxels within the explicit anatomical mask that was manually drawn to include the whole medial prefrontal cortex, caudate head and body, putamen, and pallidum. Unless otherwise stated, the statistical threshold was set at *p* < 0.05, family-wise error (FWE) corrected, *k* = 10.

First, we analyzed the data of the food gambling task to determine brain regions that correlated with estimated utility (GLM1) and amount of food (GLM2). GLM1 included five parametric regressors: (1) the utility of the sure offer; (2) the expected utility of the lottery offer weighted by subjective probability (w(*p* = 0.5)); (3) the utility of the chosen option weighted by subjective probability; (4) the utility of sure feedback; and (5) the utility prediction error of lottery feedback weighted by subjective probability at the onset of each event. The onsets of each trial per se and six motion parameters were treated as regressors of no interest. Similarly, GLM2 included five parametric regressors: (1) the amount of the sure offer; (2) the expected amount of the lottery offer weighted by subjective probability; (3) the expected amount of the chosen options weighted by subjective probability; (4) the amount of the sure feedback; and (5) the amount prediction error of the lottery feedback weighted by subjective probability at each event onset. In addition, we included the same regressors of no interest as those used in GLM1. We were able to identify the brain region whose activity correlated with the utility prediction error (from GLM1), but not with the amount prediction error (from GLM2).

Second, we analyzed the data from the monetary gambling task to determine whether the neural activity in the region whose activity was found to correlate with the utility prediction error in GLM1 was affected by the change in the width of the reward across blocks (GLM3). GLM3 included the following parametric regressors at each event onset: (1) the monetary amount of the sure offer; (2) the expected monetary amount of the lottery offer weighted by subjective probability; (3) the monetary amount of the chosen option weighted by subjective probability; (4) the monetary amount of sure feedback; (5) the amount prediction error of the lottery feedback weighted by subjective probability in the narrow block; and (6) the amount prediction error of the lottery feedback weighted by subjective probability in the wide block. In addition, we included the same regressors of no interest as those in GLM1.

We found that the neural representation of the utility prediction error (= the amount prediction error in the monetary gambling task) was not affected by the change in the width of reward across blocks. Next, we identified all the voxels representing utility prediction errors without discriminating between the narrow and the wide blocks in the monetary gambling task (GLM4). GLM4 included the same regressors as GLM3 except that the prediction error of the lottery feedback for both the narrow and the wide blocks was modeled as a single parametric regressor. By finding the overlapped region between the neural representations of the utility prediction error for money (GLM4) and food (GLM1), we determined the brain region that represents the common currency of utility. This was performed using MarsBaR (http://marsbar.sourceforge.net). All the regressors were not standardized.

### Region-of-interest (ROI) analyses

We used the rfxplot toolbox (http://rfxplot.sourceforge.net/) (Gläscher, 2009) to extract PSC in regions of interest. For the region whose activity correlated with utility prediction errors in the food gambling task, the PSCs evoked by utility prediction errors per ¥1 were compared between the narrow and the wide blocks using the classical paired t-test and the Bayesian paired t-test with default effect size priors (Cauchy scale 2^0.5^/2) (Keysers et al. 2020). Bayesian t-test results were reported using the one-tailed Bayes factor BF_+0_ that represents P(data| H_+_: μ_PSCnarrow_ > μ_PSCwide_)/P(data| H_0_: μ_PSCnarrow_ = μ_PSCwide_), where μ_PSCnarrow_ and μ_PSCwide_ indicated the mean of PSCs from the narrow and the wide blocks, respectively. Effect size estimates were reported as median posterior Cohen’s δ with 95% CI using an H_1_ without restriction of μ_PSCnarrow_ ≥ μ_PSCwide_ in order not to bias estimates in the expected direction.

We employed multiple regression analyses to evaluate the relationship between the PSCs of the “utility region” (Δu_neural_) and each participant’s household income:

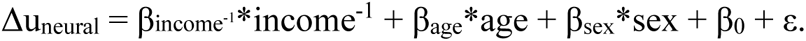

All the variables were standardized and a dummy variable was used for sex (male = 0, female = 1).

Δu_neural_ were compared between the “poor” and the “rich” groups using the classical t-test and Bayesian t-test with default effect size priors (Cauchy scale 2^0.5^/2). Bayesian t-test results were reported using the one-tailed Bayes factor BF_+0_ that represents P(data| H_+_: μ_PSCpoor_ > μ_PSCrich_)/P(data| H_0_: μ_PSCpoor_ = μ_PSCrich_), where μ_PSCpoor_ and μ_PSCrich_ indicate the mean of PSCs of the “poor” and the “rich” groups, respectively. Effect size estimates were reported as median posterior Cohen’s δ with 95% CI using an H_1_ without restriction of μ_PSCpoor_ ≥ μ_PSCrich_ in order not to bias estimates in the expected direction.

### Impartial spectator experiment: participants

Fifteen healthy student volunteers living with their parents participated in this experiment. All the participants were native Japanese speakers recruited from the same university pool as the MRI participants. None of the participants had participated in the gambling experiment or had a history of neurological or psychiatric diseases, and all the participants provided their written informed consent prior to participation. Two participants did not pass the instructional manipulation check (Oppenheimer et al. 2009) for the task in the initial instruction and were excluded from analyses, resulting in a final sample of 13 (7 female and 6 male participants; age: 20.85 ± 1.14 years, range: 19–23 years). The study was approved by the ethics committee of Tamagawa University in accordance with the ethical standards laid down in the 1964 Declaration of Helsinki and its later amendments.

### Procedures

Participants completed the impartial spectator task outside the MRI scanner.

### Impartial spectator task

First, each participant was shown a histogram of the household income distribution of the participants in the gambling experiment and told that “The amount of money you earn will depend on whether you perform the task seriously.” Next, they answered the first question: “The sum of the pleasure of the participants in the lower half of household income distribution who each received ¥500 should be balanced by the sum of the pleasure of the participants in the upper half of the household income distribution who each received ¥X_500_1_. How much is ¥X_500_1_?” In the second and third questions, ¥400 and ¥600 were used as the amounts of money that the “poorer” participants received (X_400_, X_600_). The order of ¥400 and ¥600 was counterbalanced across the participants. In the fourth question, ¥500 was once again used as the amount of money that the “poorer” participants received (X_500_2_). Participants whose responses in the first and forth trial were the same earned a bonus of ¥500.

We recruited independent participants for the impartial spectator task from the same university student pool as the MRI participants and were incentivized to perform the task seriously. Two conditions for the impartial spectators were proposed: (1) the impartiality requirement, where spectators should be third parties whose interests are not affected by their judgements about the observed; and (2) the empathy requirement, where spectators should have sufficient information and motivation to infer the pleasure of the observed (Harsanyi 1977; Smith 1759). Our participants were considered impartial and empathetic by the MRI participants, which matched the requirements for impartial spectators originally proposed by Adam Smith (Smith 1759; Harsanyi 1977).

### Validation of the interpersonal comparison of utility

The impartial spectators’ estimates of the utility difference ratio between the “poor” and the “rich” groups were calculated as X_ratio_ = (X_500_1_/500 + X_400_/400 + X_600_/600 + X_500_2_/500)/4. The impartial spectators’ estimated values of the total (or mean) utility of the “poor” and the “rich” groups were assumed to follow normal distributions with a common standard deviation (σ_IS_). Subsequently, X_ratio_ was assumed to be generated by the following formula:

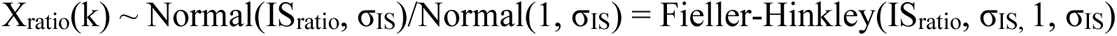

where IS_ratio_ is the mean of the estimated utility of the “poor” group when the mean of the estimated utility of the “rich” group is one.

To check whether the impartial spectators’ estimation and neural signals coincided, IS_ratio_ and the Δu_neural_ ratio of the “poor” to the “rich” group (PSC_ratio_) were compared using a Bayesian model with default effect size priors (Cauchy scale 2^0.5^/2). The results were reported as BF_10_ that represents P(data| H_1_: PSC_ratio_ ≠ IS_ratio_)/P(data| H_0_: PSC_ratio_ = IS_ratio_). Effect size estimates were also reported as median posterior Cohen’s δ with 95% CI. The model is shown in Supplementary Figure 5.

The details of the model are as follows. We first calculated the mean and standard deviation of the obtained Δu_neural_ data: 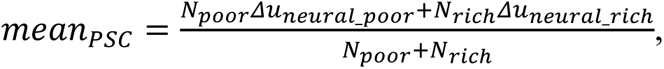 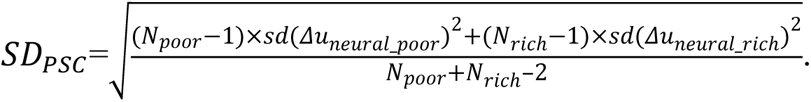We used a normal approximation of a Fieller-Hinkley distribution (Díaz-Francés and Rubio 2013) to calculate the standard deviation of Δu_neural_ ratio as 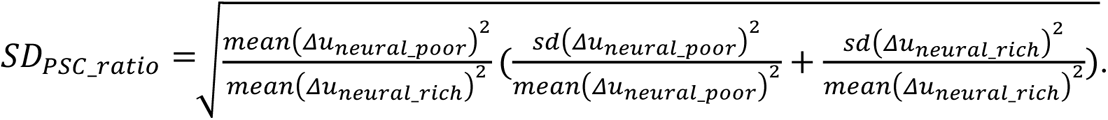 standard deviation of the ratios of the Δu_neural_ and impartial spectator data as 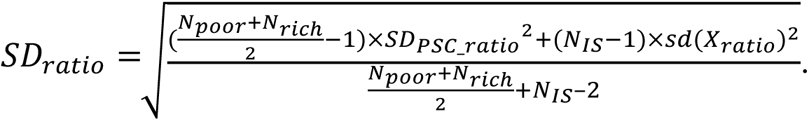 sample mean and the unbiased sample variance, respectively.

We used the following priors for the effect size, mean, and standard deviation of Δu_neural_, the effect size of the difference between Δu_neural_ data and impartial spectator data, and standard deviation of the impartial spectators’ intuition.

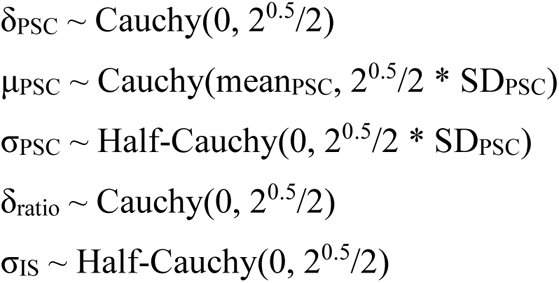

Mean difference of Δu_neural_ between the “poor” and the “rich” group (α_PSC_), PSC_ratio_, and IS_ratio_ were defined as follows.

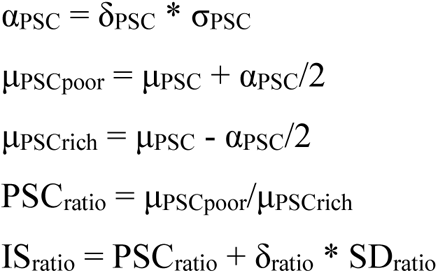

The last equation is based on the definition of Cohen’s δ (δ_ratio_ = (IS_ratio_ - PSC_ratio_)/SD_ratio_). The data were assumed to be generated using the following formula. The Δu_neural_ data were assumed to follow a normal distribution.

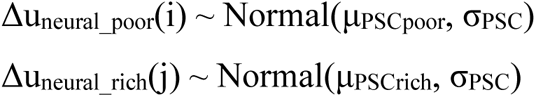

Then, X_ratio_ was assumed to be generated by the following formula:

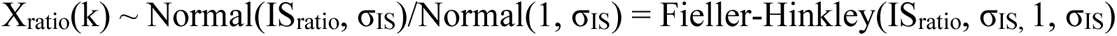

### Application of the interpersonal comparison of utility

The policy that distributes ¥x to each participant in the “poor” group and ¥y to each participant in the “rich” group was considered from a social planner’s perspective. Here, a utilitarian social welfare function was considered, because it is the most common when utility differences are interpersonally comparable (d’Aspremont and Gevers 1977; Roberts 1980).

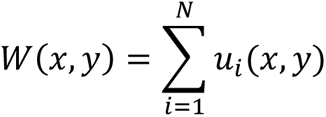

The increase in social welfare generated by the policy for moderate amounts of money was expressed as follows:

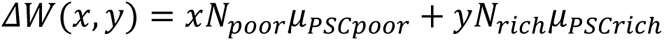

Given posterior of P(μ_PSCpoor_) and P(μ_PSCrich_) in the Bayesian t-test model that has H_1_ without restriction of μ_PSCpoor_ ≥ μ_PSCrich_, the expected social welfare increase was expressed as follows.

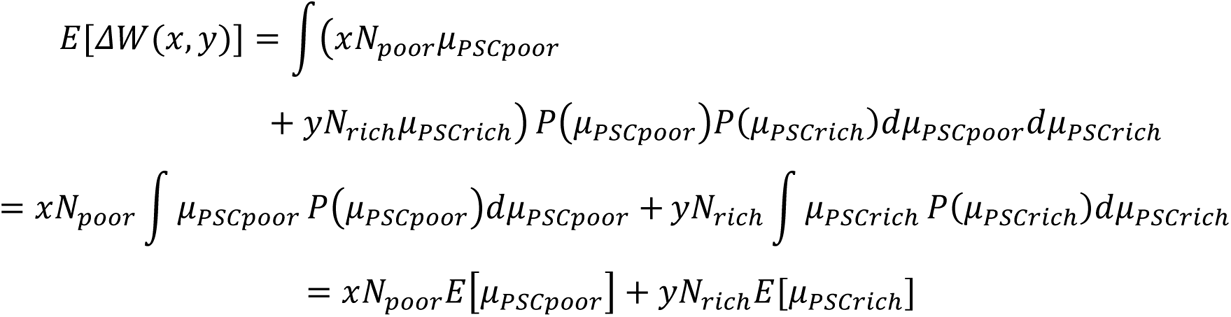

The expected social welfare could be maximized by choosing the option with maximum E[ΔW]. This equation was used to determine the optimal decision rule for the problem of choosing between a policy that distributes ¥k to all the participants and a policy that distributes ¥(k + ρ) to each participant in the “poor” group. If the two policies generate the same increase in social welfare, the following equation holds:

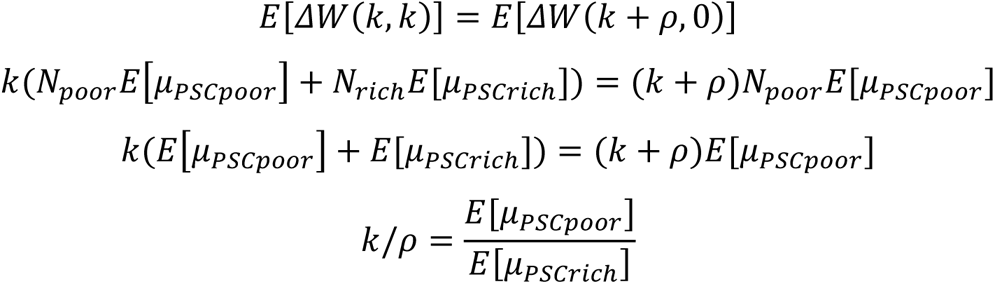

Therefore, the optimal decision rule for expected utilitarian social welfare maximizer, is

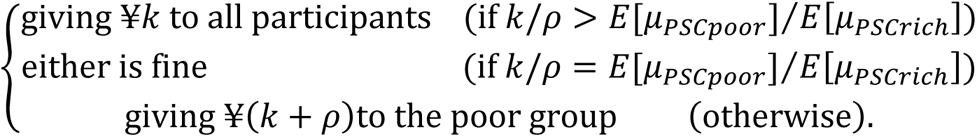

This decision rule was applied to choose between a policy that distributes ¥1,000 to all the participants and a policy that only distributes ¥1,500 to participants in the “poor” group. Based on the neural data, we allocated an additional ¥1,500 to all the “poor” participants who agreed to receive additional compensation (n = 13).

### ABCD study data analysis: participants

Participants were selected from the ABCD study, which was designed to recruit a representative cohort of children in the United States population (http://abcdstudy. org/). The ABCD study was approved by the institutional review board (IRB) of each study site, with centralized IRB approval from the University of California, San Diego. Informed consent was obtained from all the parents and children. Initially, we considered a sample of 6,803 children (aged 9–10 years) who had minimally processed fMRI data available for both monetary incentive delay (MID) task runs and who had passed the ABCD MRI quality checks for both runs. Participants with technical problems with their MRI data processing (n = 33) and those with no household income data (n = 549) were excluded from the analysis. To control for family influence, we randomly selected one participant from each family, which resulted in the exclusion of 849 participants.

The final sample comprised 5,372 children. Household income data was collected using a 10-level scale: 1 = less than $5,000; 2 = $5,000–$11,999; 3 = $12,000–$15,999; 4 = $16,000–$24,999; 5 = $25,000–$34,999; 6 = $35,000–$49,999; 7 = $50,000–$74,999; 8 = $75,000–$99,999; 9 = $100,000–$199,999; and 10 = $200,000 and greater. For the analysis, each income category was represented by its midpoint value. The highest category (10) was represented by $250,000.

### The MID task

Casey et al.’s (2018) study offers a comprehensive overview of the MID task and the associated fMRI data collection methods used in the ABCD study. In the task, each trial began with an incentive cue, which indicated whether the participants had an opportunity to win money (reward trial), face a potential monetary loss (loss trial), or experience no monetary stakes (neutral trial). After the cue, the participants responded to a target as quickly as possible, with the task adjusted to maintain an approximate 60% success rate. Timely responses resulted in positive feedback (winning money or avoiding a loss), whereas late responses led to negative feedback (failing to win or losing money). As per the standard MID procedure (Knutson et al. 2001), trials alternated between small ($0.20) and large ($5.00) incentives for both reward and loss trials. The task was divided into 40 reward trials, 40 loss trials, and 20 neutral trials, which were distributed evenly across the two fMRI runs.

### MRI data acquisition

Functional imaging was conducted using three different 3T MRI scanner platforms (Siemens Prisma, General Electric 750, and Phillips) at 21 sites. Gradient echo T2*-weighted EPI were acquired using blood oxygenation level-dependent contrasts. Sixty EPI slices were acquired in each volume (slice thickness, 2.4 mm; repetition time, 800 ms; echo time, 30 ms; flip angle, 52°; field of view, 216 mm^2^; matrix, 90 × 90; multiband factor = 6). Data were acquired for each participant in two scanning sessions for the MID task. Initial dummy volumes for each session were discarded according to the length of the dummy events for each MRI model to allow for T1 equilibrium effects.

### Preprocessing of functional images

The minimally processed neuroimaging data were downloaded, and the following preprocessing procedures were performed using SPM12 software (Wellcome Department of Imaging Neuroscience, London, UK, https://www.fil.ion.ucl.ac.uk/spm/) implemented in MATLAB: adjustment of acquisition timing across slices, spatial normalization to the standard MNI template, and smoothing using a Gaussian kernel with a full width at half maximum of 8 mm.

### Functional MRI data analysis

We adopted a conventional approach using SPM12 to analyze fMRI data. A set of regressors was generated by convolving the canonical hemodynamic response function provided by SPM12 with a series of epochs (cue presentation, feedback). A voxel-by-voxel multiple regression analysis was applied to the preprocessed images for each participant. We analysed the data from the MID task to calculate beta maps for each event. The GLM at each event onset included two parametric regressors: (1) the amount of the cue and (2) the amount of the prediction error of feedback. Onsets of each trial per se and six motion parameters were treated as regressors of no interest. The MarsBaR toolbox was used to extract PSC at regions of interest determined in the gambling experiment. The PSC per $1 (Δu_neural_) were calculated for each participant.

To evaluate the relationship between the Δu_neural_ and each participant’s household income, we employed a linear mixed model using the statsmodels package in Python. The model was specified as follows:

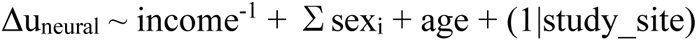

where income^-1^ is the reciprocal of household income, and sex_i_ is the dummy variable for the sex categories. The reciprocal of household income, sex, and age are fixed effects, and study_site is included as a random effect to account for site-specific variations. Household income was converted from categorical values to actual income amounts using the midpoint value of each income category. Continuous variables (Δu_neural_, age and the reciprocal of household income) were standardized.

To estimate the elasticity of the utility function with respect to income, we employed a nonlinear model using an isoelastic utility function. The model was specified as:

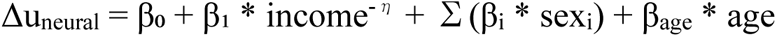

where η represents the non-negative elasticity parameter of the isoelastic utility function. We used a bootstrap approach with 1,000 iterations to estimate the parameters and their confidence intervals, resampling with replacements from the original dataset for each iteration. The model was fitted using Python’s SciPy optimization function “minimize” with the SLSQP method.

## Data availability

The PSC data and impartial spectator behavioral data will be available from the GitHub repository. The anatomical mask and statistical fMRI maps associated with this study are available at the NeuroVault repository (https://neurovault.org/collections/YVHREMBI/). The original ABCD data are available with permissions from the NIMH data archive (https://nda.nih.gov/abcd).

## Code availability

The custom code supporting the results of this study will be available from the GitHub repository.

## Acknowledgments

We would like to thank Reiko Gotoh, Ralph Adolphs, Mikihiko Wada, Masanori Kataoka, Shu Ishida, Kou Murayama, Tatsuya Kameda, and Tsuguo Mogami for their valued discussions. This study was partially supported by a Grant-in-Aid for JSPS Research Fellow Number JP20J01726 and Moonshot R&D JPMJMS2294 (to Ka.M.), JSPS KAKENHI Grant Numbers 17H05929 (to K. I.), JSPS KAKENHI Grant Numbers JP18H05085 and JP19H04885, and AMED Grant Numbers JP18dm0207008 and JP21dm0307001 (to Ke.M). .

## Author contributions

Ka.M. conceived the idea, and Ka.M., K.I. and Ke.M jointly designed the study. Ka.M., Y.Y., and Ke.M. created the experimental materials. Ka.M., K.I., and Ke.M. performed the research. Ka.M. analyzed the data. Ka.M., K.I., and Ke.M. jointly wrote the paper. Ka.M., K.I., Y.Y., and Ke.M. provided critical comments.

## Competing interests

The authors declare no competing interests.

**Correspondence and requests for materials** should be addressed to Ka. M.. and Ke.M.

## Supplementary information

### Section 1: detailed introduction of utility concepts

Bentham’s (1789) original understanding of utility (psychological utility) was psychological pleasure. For example, if one person’s pleasure when eating an apple is greater than when eating an orange, then their psychological utility for the apple is greater than their utility for the orange. Until the beginning of the 20th century, economists regarded utility in this sense as interpersonally comparable. It is often pointed out that, in everyday life, ordinary people make interpersonal comparisons of utility with relative ease and apparent success (Davidson 2004; Rossi 2014). Most people would think, “If a poor person and a rich person receive the same amount of money, the poor person will be more pleased than the rich person.”

However, psychological utility cannot be directly measured by behavior. Along with the rise of logical positivism and behaviorism, a new concept of utility (decision utility) that could be operationally defined by an agent’s behavior emerged as a surrogate for psychological utility. As agents are usually thought to prefer an option with greater psychological utility, a function *u,* such that *u*(preferred option) is greater than *u*(non-preferred option), came to be called the utility function (Kreps 2012; Moscati 2018). In this concept, if an agent prefers an apple to an orange, they have greater decision utility for an apple than for an orange, and this is measurable by an agent’s choice behavior. Thus, decision utility was considered an ordinal scale, because only the relationship “prefer to” was available (ordinal utility) (Figure 1A).

Von Neumann and Morgenstern attempted to turn the concept of utility into an interval scale by considering the history of the development of the temperature scale (Arrow 1963; Von Neumann and Morgenstern 2007). When only the relationship “warmer than” was known, temperature was considered an ordinal scale. Temperature became an interval scale (e.g., Celsius and Fahrenheit) through examinations of the expansion of fluids, such as mercury, at the beginning in the 17th century. The initial validation of these attempts was based on a coherence between such scales and human sensation (Chang 2004; Tal 2015), namely, obvious observations of cold and heat in winter and summer. Temperature scales that are based on the expansion of fluids have two free parameters for unit width (α) and zero point (β), and are unique up to positive linear transformation (x’ = αx + β, α > 0) mathematically. The Celsius and Fahrenheit scales have different unit intervals and zero points, and they are commensurable with a linear transformation. Subsequent developments in thermodynamics made temperature a ratio scale (Kelvin scale).

Von Neumann and Morgenstern developed the concept of the utility function on lotteries as an interval scale (Krantz et al. 1971; Von Neumann and Morgenstern 2007) (cardinal utility) (Figure 1B, C). Here, we assume an agent’s preference order is apple, banana, and orange. In this concept, if an agent is indifferent to a banana and a 50/50 gamble of an apple and an orange, the difference in utility between an apple and a banana is equal to the difference in utility between a banana and an orange (Figure 1B, C). Cardinal utility is also unique up to linear transformation, and any scale based on cardinal utility is commensurable with a linear transformation. Thus, Von Neumann and Morgenstern succeeded in expressing utility as interval scale. Later studies on choice probability, decision time, and neural signals supported this idea (Abdellaoui et al. 2007; Luce 1959; Ratcliff et al. 2016; Stauffer et al. 2014; Stott 2006; Waldner 1972). A study has also shown that expected utility weighted by subjective probability has a linear relationship with psychological utility (Abdellaoui et al. 2007), suggesting the replacement of psychological utility with expected utility weighted by subjective probability (Dietrich and List 2016; Okasha 2016).

However, this type of utility is based only on individual choice behavior; only the relative position of the utility for an option between the best and worst options on an arbitrary linear scale is empirically measurable (Figure 1B, C, E, F). In other words, the information of utility as interval scale is fully preserved even after a linear transformation with the parameters for different unit width (α_i_) and for different zero point (β_i_) among individuals (u_i_’ = α_i_u_i_ + β_i_, where *i* is for individuals). A popular transformation is the expression of a scale from 0 for the worst option to 1 for the best option (0–1 rescaling). This indicates that decision utilities are comparable by definition only *intra*personally, not *inter*personally. Note that this limitation applies to ordinal utility as well.

Thus, after the 1930’s, interpersonal comparisons of utility based on lay people’s intuition came to be regarded as neither objective nor scientific and remained a taboo for nearly a century since the concept of decision utility was popularized (Robbins 1932, 1938). This is a serious problem caused by the replacement of psychological utility with decision utility. Using decision utility alone, the utility of a slice of bread cannot be compared even between a starving person and a billionaire (Sen 2018).

### Section 2: introduction to neuroscientific studies

A natural and scientific extension of decision utility would be to use neural data (Narens and Skyrms 2020). First, any economic choice behavior that reveals decision utility consists of voluntary muscle movements, which are generated by signals that originate in the brain. Second, it is natural to assume that economic choice behavior that reveals decision utility is determined by psychological utility. Furthermore, mental states— including psychological utility—are realized by brain states. Therefore, psychological utility should be reflected in the activities of the responsible brain areas.

Neuroscientific studies using rodents, monkeys, and humans have repeatedly shown that damage to specific brain regions, such as the ventral/medial prefrontal cortices, the ventral striatum, and the midbrain dopaminergic nuclei causes impairments in choice behaviors (Butter et al. 1969; Camille et al. 2011; Cardinal et al. 2001; Frank et al. 2004). In addition, a recent study reported that electrical stimulation of the ventral prefrontal cortex causes changes in monkey choice behaviors (Ballesta et al. 2020). Functional neuroimaging studies of human participants have shown that the activities of these regions of the brain represent the value of various rewards (e.g., food, money, and social praise) on a single common scale (Bartra et al. 2013; Glimcher and Fehr 2013; Izuma et al. 2008; Levy and Glimcher 2012). These areas of the brain are associated with rewards or value systems; moreover, reward-related regions are less responsive to food when not hungry (Cassidy and Tong 2017; Gottfried et al. 2003) and less responsive to monetary rewards when the reward is delayed ( Kable and Glimcher 2007; McClure et al. 2004), which suggests that the activities of these regions represent the subjective aspects of the value of rewards. Therefore, utility information is expected to be found in reward-related activities.

One study using monkey subjects showed that neuronal activity in the midbrain dopaminergic nuclei, which project to the reward-related regions, encoded utility difference (Δu) (Stauffe et al. 2014). If we find a utility representation in human brains, it may allow us to objectively measure utility differences between options associated with individual mental states, in contrast to measurement based on choice behaviour alone (Figure 1D). The information of such a utility representation should be preserved after a linear transformation with the parameters for common unit width (α) and for different zero points (β_i_) among individuals (u_i_’ = αu_i_ + β_i_, where *i* is for individuals).

## Supplementary figures

**Supplementary Figure 1.**
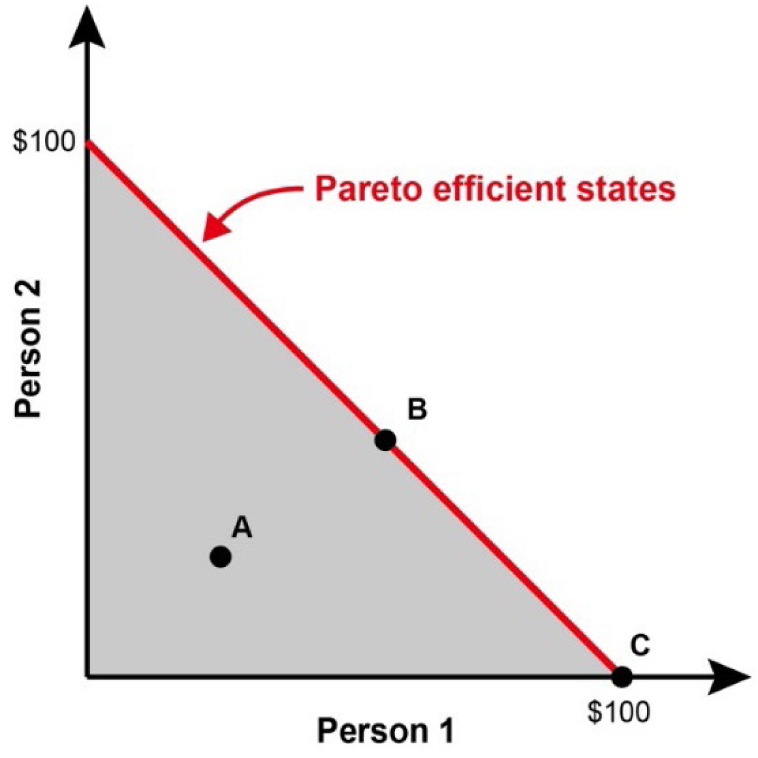
Pareto efficiency. Consider the situation in which Persons 1 and 2 split $100. The feasible states are shown in a gray area. Under the assumption that each person’s utility is an increasing function of money, Point A is not Pareto efficient, because both persons’ share can increase from point A to B. Point B is Pareto efficient because no one’s share can increase without decreasing the other’s share. Any point on the red line is in a Pareto efficient state; however, Pareto efficient states do not exclude highly unequal points such as point C.

**Supplementary Figure 2.**
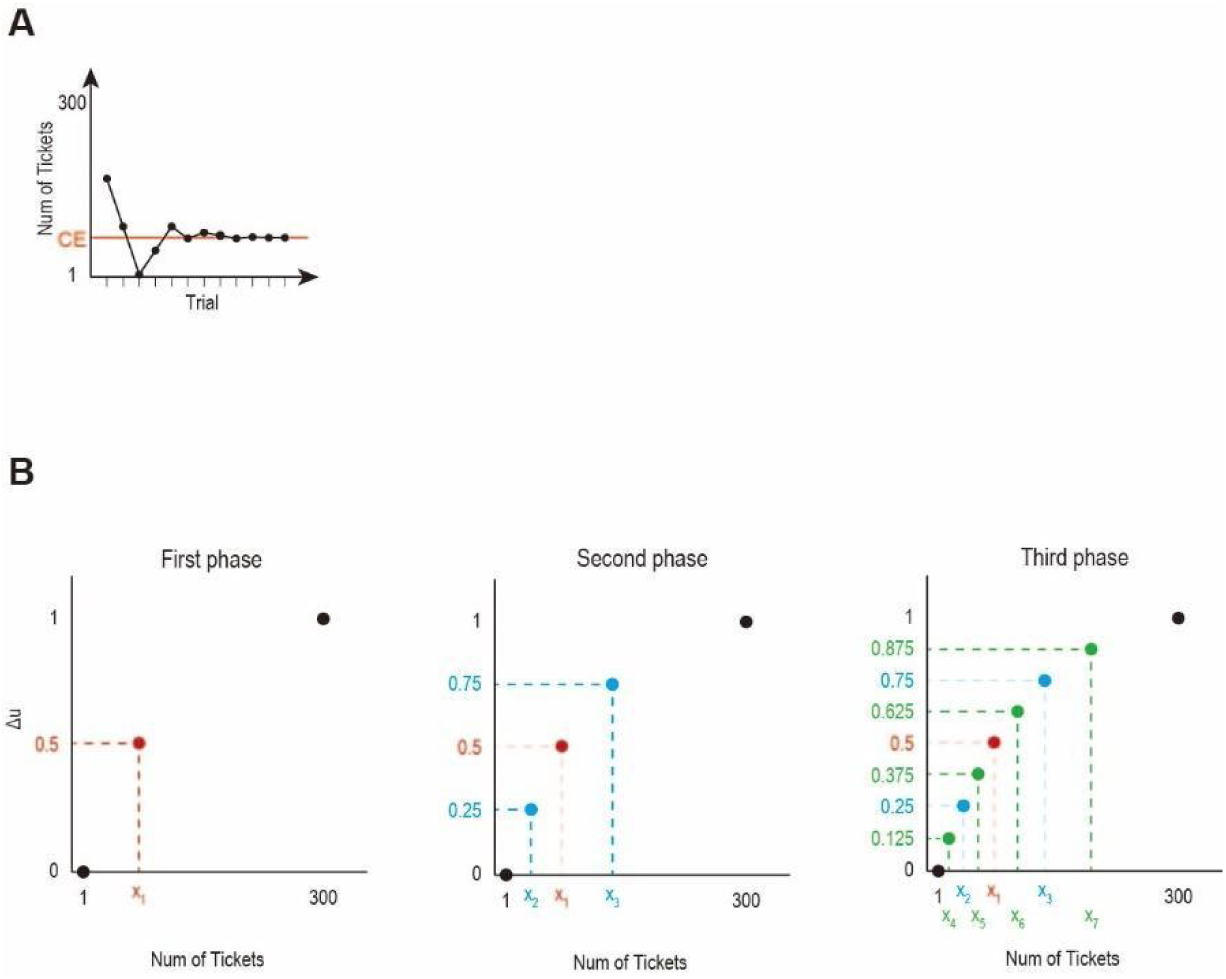
Measuring utility function. (**A**) Measuring CEs using the PEST procedure. The black trace shows the number of tickets for a sure option compared with the 50/50 gamble between 1 and 300 tickets in the food gambling task. The lottery loss number (= 1) and lottery win number (= 300) remained unchanged throughout a PEST sequence. The sure number was adjusted in subsequent trials based on the participant’s choice between the sure offer and lottery (see *Behavioral analyses: measuring CEs*). The procedure lasted until convergence. (**B**) Iterative fractile method for measuring utility in the food gambling task. First, the CE of the 50/50 gamble between 1 and 300 tickets (x1) was measured by using PEST (red point). Then, x_1_ was used as an outcome to construct two new gambles (1 ticket, *p* = 0.5 and x_1_ tickets, *p* = 0.5; x_1_ tickets, *p* = 0.5 and 300 tickets, *p* = 0.5) and measure their CEs (x_2_ and x_3_, light blue points). Finally, the CEs of the 50/50 gamble between 1 and x_2_ tickets (x_4_), between x_2_ and x_1_ tickets (x_5_), between x_1_ and x_3_ tickets (x_6_), and between x_3_ and 300 tickets (x_7_) were measured (light green points). The figure illustrates the case where w(*p* = 0.5) = 0.5. If w(*p* = 0.5) = 0.4, the Δu(x_1_) is 0.4 rather than 0.5 and so on; w(*p* = 0.5) was determined by the probability weighting task (see *Methods*).

**Supplementary Figure 3.**
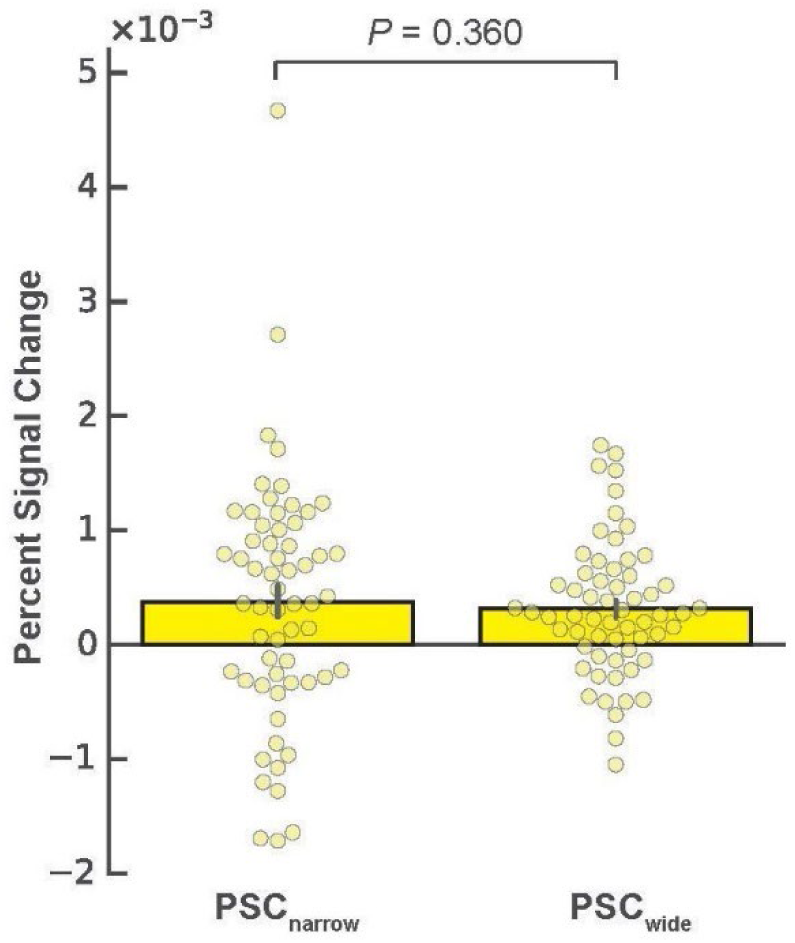
Neural signals for the utility prediction errors in two reward-range conditions. The chart shows the activity of the brain region representing utility prediction errors in the food gambling task. There was moderate evidence that the mean percent signal changes for utility prediction errors per 1 yen were not rescaled between the narrow and the wide blocks in the monetary gambling task (t(59) = 0.36, *P* = 0.360 one-sided, BF_+0_ = 0.193, median posterior δ = 0.044 with 95% CI [-0.201, 0.290]). Error bars indicate SEM across participants.

**Supplementary Figure 4.**
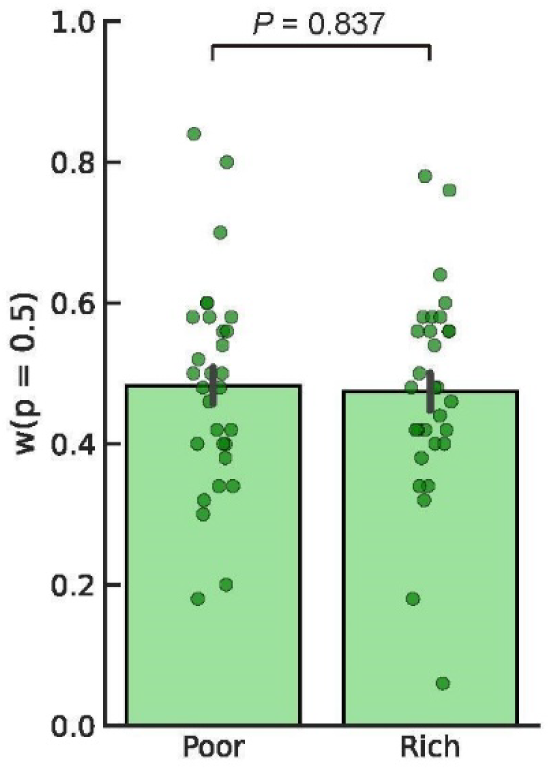
Probability weighting of the “poor” and the “rich” groups of participants. The chart shows the subjective probabilities at *p* = 0.5 (w(*p* = 0.5)) of the “poor” and the “rich” groups of participants in the probability weighting task. The values did not differ between the two groups with moderate evidence (t(58) = 0.207, *P* = 0.837 two-sided, BF_10_ = 0.268, median posterior δ = 0.044 with 95% CI [-0.418, 0.510]). Error bars indicate SEM across group members.

**Supplementary Figure 5.**
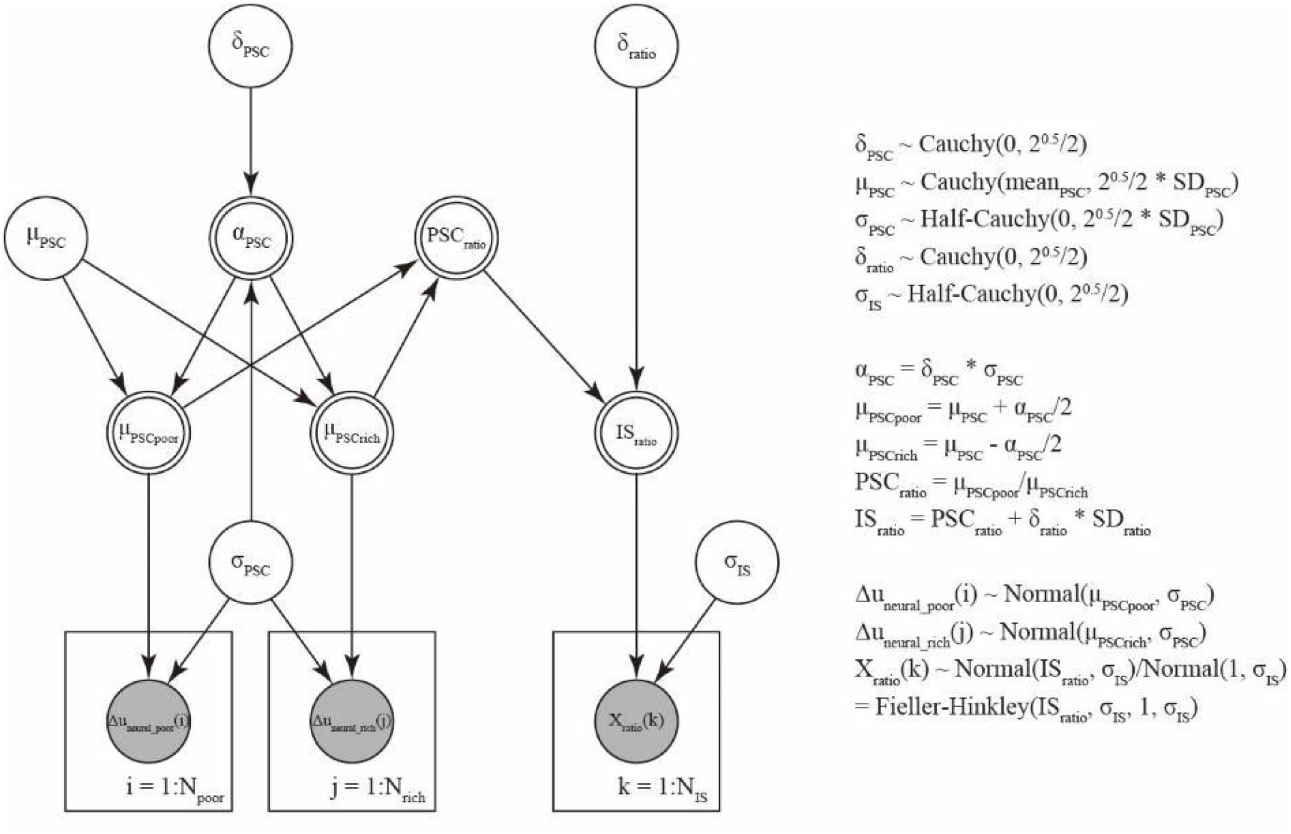
Graphical model to assess coincidence between neural signals and impartial spectators’ estimations. The shaded and unshaded nodes represent unobserved and observed variables, respectively. Single-bordered and double-bordered nodes represent probabilistic and deterministic variables, respectively.

